# MOZ and HBO1 Histone Acetyltransferase Complexes Are Molecular Dependencies and Therapeutic Targets in *NUP98*-Rearranged Acute Myeloid Leukemia

**DOI:** 10.1101/2024.12.02.624182

**Authors:** Nicole L. Michmerhuizen, Emily B. Heikamp, Ilaria Iacobucci, Masayuki Umeda, Bright Arthur, Vibhor Mishra, Danika Di Giacomo, Ryan Hiltenbrand, Qingsong Gao, Sandi Radko-Juettner, Josi Lott, Cynthia Martucci, Varsha Subramanyam, Charlie Hatton, Pradyuamna Baviskar, Pablo Portola, Aurelie Claquin, Bappaditya Chandra, David W. Baggett, Ali Khalighifar, Hongling Huang, Peipei Zhou, Lingyun Long, Hao Shi, Yu Sun, Evangelia K. Papachristou, Chandra Sekhar Reddy Chilamakuri, Francisca N. de L. Vitorino, Joanna M. Gongora, Laura Janke, Alex Kentsis, Clive S. D’Santos, Benjamin A. Garcia, Richard W. Kriwacki, Hongbo Chi, Jeffery M. Klco, Scott A. Armstrong, Charles G. Mullighan

## Abstract

NUP98 fusion oncoproteins (FOs) are a hallmark of childhood acute myeloid leukemia (AML) and drive leukemogenesis through liquid-liquid phase separation-mediated nuclear condensate formation. However, the composition and consequences of NUP98 FO-associated condensates are incompletely understood. Here we show that MYST family histone acetyltransferase (HAT) complex proteins including MOZ/KAT6A, HBO1/KAT7, and the common MOZ/HBO1 complex subunit BRPF1 associate with NUP98 FOs on chromatin and within condensates. MYST HATs are molecular dependencies in *NUP98*-rearranged (*NUP98*-r) leukemia, and genetic inactivation or pharmacologic inhibition of *Moz* and *Hbo1* impairs *NUP98*-r cell fitness. MOZ/HBO1 inhibition decreased global H3K23ac levels, displaced NUP98::HOXA9 from chromatin at the *Meis1* locus, and led to myeloid cell differentiation. Additionally, MOZ/HBO1 inhibition decreased leukemic burden in multiple *NUP98*-r leukemia xenograft mouse models, synergized with Menin inhibitor treatment, and was efficacious in Menin inhibitor-resistant cells. In summary, we show that MYST family HATs are therapeutically actionable dependencies in *NUP98*-r AML.

**SIGNIFICANCE STATEMENT:** MOZ and HBO1 associate with NUP98 fusion oncoprotein condensates to drive leukemogenesis. Inhibition of their histone acetyltransferase activity is an effective therapeutic strategy in *NUP98*-rearranged leukemias, including those resistant to Menin inhibition. Moreover, combined MOZ/HBO1 and Menin inhibition is synergistic, supporting clinical translation to improve outcomes of NUP98 FO-driven leukemias.

## INTRODUCTION

Acute myeloid leukemia (AML) is the second most common form of pediatric leukemia (1), and chromosomal rearrangements are detected in most pediatric AML patients. NUP98 fusion oncoproteins (FOs) are observed in ∼5% of pediatric AML, and at higher rates in monocytic, megakaryoblastic, and erythroid AML (2–6). Approximately 70% of children with *NUP98*-rearranged (*NUP98*-r) AML experience disease relapse after standard therapy, constituting one of the most common drivers of relapsed disease in pediatric AML (2–4,6–9). More effective therapeutic options for *NUP98-*r AML are needed, and the development of more improved therapies requires improved mechanistic understanding of NUP98 FO-driven leukemogenesis.

NUP98 is a component of the nuclear pore complex, which mediates the transport of macromolecules between the cytoplasm and nucleus (10,11). NUP98 FOs retain the N-terminal Gle2-binding-sequence (GLEBS) domain and intrinsically disordered glycine-leucine-phenylalanine-glycine (GLFG) repeats of NUP98 (11). More than 30 different NUP98 fusion partners have been identified, many of which have a DNA-binding homeodomain (12). The majority of fusion partners lacking this domain, including the most prevalent partners, KDM5A and NSD1, harbor domains involved in writing or reading histone modifications. Chromatin remodeling by NUP98 FOs drives self-renewal and augments proliferation (13,14). NUP98 FOs upregulate targets including *HOX* gene clusters and *MEIS1*, which are otherwise silenced during hematopoietic differentiation (13–15).

NUP98 FOs localize in nuclear foci or puncta, and recent work from our group and others demonstrated that these puncta function as transcriptional condensates via liquid-liquid phase separation (16–18). The intrinsically disordered region of NUP98, which is present in all NUP98 FOs, forms gel-like condensates in vitro (19). Through a combination of homotypic and heterotypic interactions, NUP98 FO nuclear puncta orchestrate cell transformation and leukemic gene expression patterns (17,18). Previous studies have identified NUP98 FO interactors including proteins involved in nuclear export (i.e. XPO1) and ribosome biogenesis (i.e. DDX24) (16,20). Importantly, transcriptional regulators also interact with NUP98 FOs, including members of the lysine methyltransferase 2A (KMT2A/MLL)-Menin and WDR-SET-COMPASS chromatin-modifying complexes (21,22). FO-associated puncta containing XPO1 and KMT2A associate with chromatin at critical developmental loci such as the *Hoxa* gene cluster in mouse embryonic stem cells (23). Furthermore, disruption of the KMT2A-Menin complex via treatment with small molecule inhibitors of the Menin-KMT2A protein-protein inhibition (i.e. VTP-50469, SNDX-5613/Revumenib) displaced NUP98 FO from chromatin and extended survival in mice bearing *NUP98*-r patient-derived xenografts (PDXs) (24). Despite these insights on the functional role of FO in transcriptional condensates and the therapeutic potential of disrupting FO-associated chromatin complexes, it remains unknown which chromatin complexes are present in puncta, how they contribute to NUP98 FO-driven phenotypes, and whether they might represent therapeutic dependencies.

Here, we employ integrated proteomic and functional genomic approaches using mouse and human models of *NUP98-*r leukemia, which nominated and demonstrated the role of MYST histone acetyltransferase (HAT) complexes in NUP98 FO-driven leukemias. Our results point to a critical role for MYST family HAT complexes that acetylate lysine residues of the tail of histone H3, namely lysine acetyltransferase 6A (KAT6A/MOZ) and lysine acetyltransferase 7 (KAT7/HBO1). We show that members of these MYST family HAT complexes interact with various NUP98 FOs and play critical roles in *NUP98*-r leukemia cell proliferation and gene regulation. Finally, we demonstrate that small molecule inhibitors of MOZ and HBO1 are effective in preclinical models of *NUP98*-r leukemia, and that MOZ/HBO1 and Menin inhibition are synergistic, suggesting their potential therapeutic use in these high-risk leukemias.

## RESULTS

### Mouse leukemia models recapitulate NUP98-r disease in vitro and in vivo

Multiple human and mouse models for studying NUP98 FOs in the context of different C-terminal translocation partners with various epigenetic functionalities have been generated using viral and conditional knockin approaches. These models include a widely used genetically engineered mouse model of NUP98::HOXD13, which models myelodysplastic syndrome with progression to acute leukemia in some animals (12,25). However, studies systemically comparing the contribution of various C-terminal translocation partners are lacking. Of more than thirty reported *NUP98* fusion partner genes (12), most have roles in DNA binding or chromatin remodeling. Here we selected eight NUP98 FOs observed in pediatric leukemia and other hematologic malignancies. These fusions represent varied clinical characteristics and diverse fusion partners for detailed investigation (Figure 1A). We studied NUP98::KDM5A and NUP98::NSD1, the two most frequent NUP98 FOs (5,26). These two FOs harbor plant homeodomain (PHD) and Suppressor of variegation 3-9, Enhancer of zeste, and Trithorax (SET) domains, which respectively play roles in “reading” histone posttranslational modifications and in methylation of histone and non-histone targets. We also studied *NUP98::JADE2*, another PHD domain-containing fusion, which has been reported in 2 cases of pediatric myeloid neoplasms with a distinct clinical phenotype (27,28), as well as *NUP98::HOXA9* and *NUP98::PRRX1*, which both contain homeodomains and are observed in a variety of hematologic malignancies. Also included were translocations of *NUP98* with *LNP1* and *RAP1GDS1*, fusion partners lacking known DNA binding or chromatin modifying domains. *NUP98::LNP1* has been reported in AML, myelodysplastic syndrome (MDS), and T-cell acute lymphoblastic leukemia (T-ALL) (29,30), and *NUP98::RAP1GDS1* is the most common *NUP98* fusion in T-ALL (31,32). Finally, we selected *NUP98::SETBP1*, another fusion previously reported in T-ALL that contains a SET binding domain (33).

**Figure 1.**
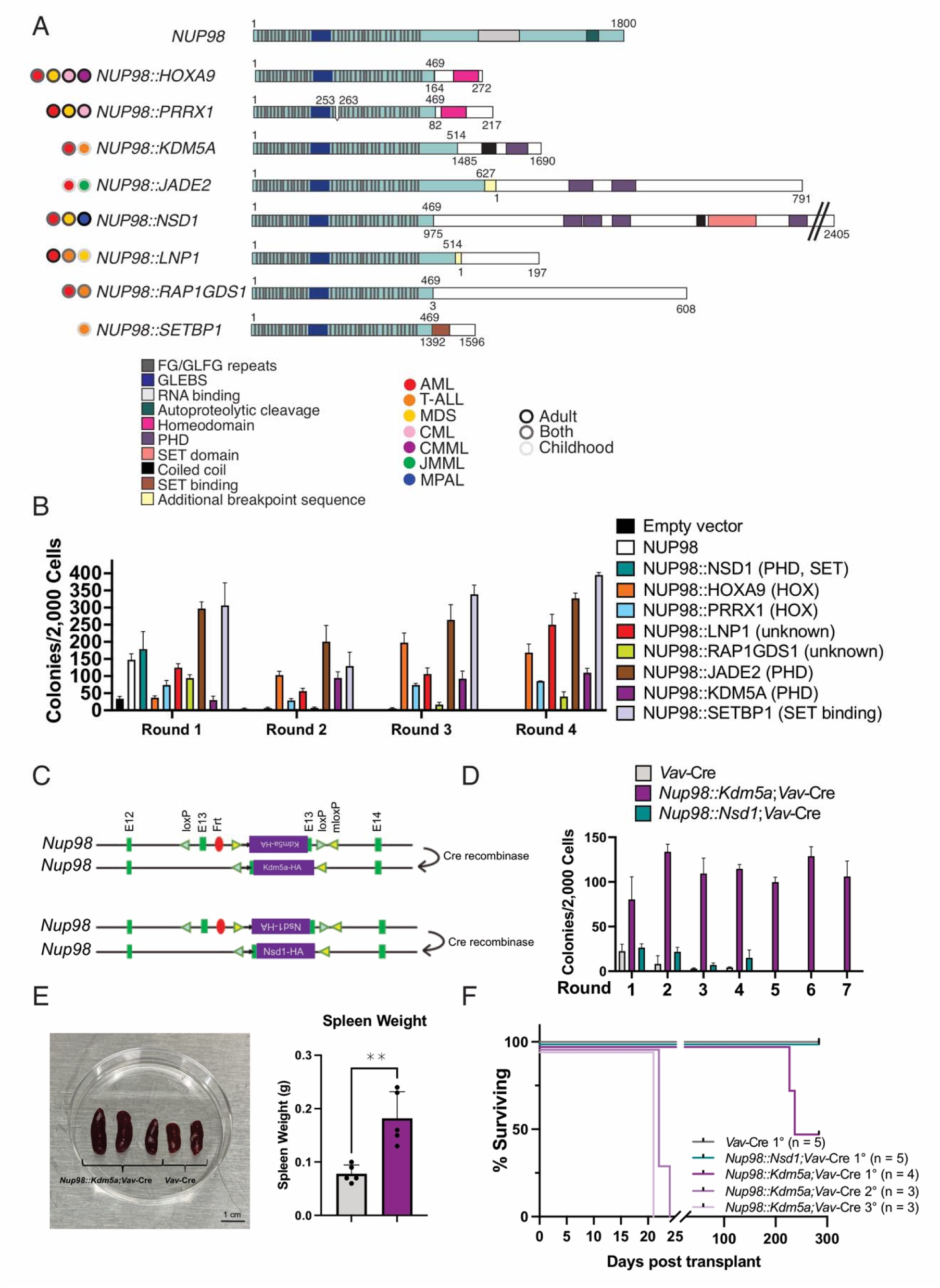
NUP98 FOs with diverse C-terminal partners transform mouse hematopoietic progenitors. A) Schematic of wildtype NUP98 and NUP98 FOs examined in this study. B) Lineage negative (lin-) hematopoietic stem and progenitor cells (HSPCs) from wildtype C57Bl/6 mice were transduced with empty vector, NUP98, or NUP98 fusion oncoprotein (FO) and colony forming unit assays were performed. C) Schematic of conditional knock in mouse models with *Nup98::Kdm5a* or *Nup98::Nsd1* fusion. D) Lin-HSPCs were isolated from *Nup98::Kdm5a;Vav* -Cre, *Nup98::Nsd1;Vav*-Cre, or wildtype *Vav*-Cre littermates, and colony forming unit assays were performed. E) Image and quantification of spleens from 3-month-old *Nup98::Kdm5a;Vav*-Cre mice or wildtype Vav-Cre littermates. ** denotes *P*< 0.01, paired t-test. F) Survival curve for mice transplanted lin-HSPCs isolated from *Nup98::Kdm5a;Vav*-Cre, *Nup98::Nsd1;Vav*-Cre, or wildtype *Vav*-Cre littermates. Spleen cells from sick mice were transplanted into secondary and tertiary recipients.

Each of these NUP98 FOs were retrovirally expressed in mouse lineage-negative (lin-) HSPCs, and colony-forming unit (CFU) assays were performed under myeloid conditions to assess their potential for cell transformation. Expression of 7 of 8 (87.5%) FOs showed enhanced clonogenic potential represented by serial replating, in contrast to cells expressing empty vector or wildtype full-length NUP98. As the sole exception, expression of NUP98::NSD1 did not lead to serial replating (Figure 1B). As we have shown previously for a subset of fusion partners (HOXA9, KDM5A, LNP1, PRRX1) (18), immunophenotyping of transformed colonies indicated expression of myeloid markers CD11b and/or Gr1 for each transforming fusion (SFig 1A).

We extended our in vitro results by generating conditional knock in mouse models of the two most common NUP98 FOs, NUP98::KDM5A and NUP98::NSD1, in which FO expression is driven by the endogenous *Nup98* promoter in the presence of Cre recombinase (Figure 1C). *Nup98::Kdm5a* and *Nup98::Nsd1* mice were crossed with *Vav*-Cre animals to express FO in the hematopoietic lineage. We performed CFU assays with lin-HSPCs from *Nup98::Kdm5a;Vav*-Cre mice, *Nup98::Nsd1;Vav*-Cre mice, or *Vav*-Cre littermates (Figure 1D). Our results recapitulated observations using retroviral overexpression of NUP98::KDM5A or NUP98::NSD1, as only NUP98::KDM5A FO expression was sufficient for cell transformation (Figure 1B, SFig 1A). We also observed increased spleen weight in 3-month-old *Nup98::Kdm5a;Vav*-Cre mice as compared to wildtype littermates of the same age (Figure 1E). Furthermore, transplantation of *Nup98::Kdm5a;Vav-Cre* lin-HSPCs into lethally irradiated congenic recipients led to the development of myeloid leukemia in 50% (2 of 4) of mice with a median latency of 232 days (Figure 1F, SFig 1D-I, Supplementary Results). We also monitored *Nup98::Kdm5a;Vav*-Cre mice to determine if they would spontaneously develop disease, and several animals developed myeloid leukemia without bone marrow transplantation (SFig 2, Supplementary Results).

### NUP98 fusion oncoproteins interact with MYST family histone acetyltransferase complexes

We investigated the protein interactome of our diverse models of NUP98 FOs (Figure 1A) using rapid immunoprecipitation mass spectrometry of endogenous proteins (RIME) combined with label free quantification, which enriches for interactions on chromatin and allows quantitative comparisons between different conditions (34,35). HEK293T cells were transiently transfected to express empty vector, HA-NUP98, or HA-NUP98 FO, and immunoprecipitation (IP) was performed using an HA antibody. Liquid chromatography-mass spectrometry (LC-MS/MS) analysis revealed known interactors of NUP98 and NUP98 FOs, including XPO1, RAE1, KMT2A, and Menin (21,36–38) (Figure 2A-B, SFig 3A). We also observed interactions with MYST family HAT complex members, including those with enzymatic (MOZ, HBO1), reader (BRPF1), and scaffolding (MEAF6) functions (Figure 2A-B). Furthermore, gene set enrichment analysis (GSEA) of NUP98::KDM5A RIME data compared to empty vector data showed enrichment of histone H3 acetyltransferase complex members (SFig 3B). NUP98::KDM5A and native NUP98 share a large portion of the interactome, whereas BRPF1, an epigenetic reader and HAT complex member, was among the most significantly enriched interactors along with NUP98::KDM5A (SFig 3C-D). Finally, we performed analogous RIME experiments for 7 additional NUP98 FOs (Figure 1A), which showed evidence of interaction between multiple NUP98 FOs and BRFP1, MEAF6, MOZ, and/or HBO1 (SFig 3E). IP-western blot analysis confirmed the interaction between HA-tagged NUP98::KDM5A, NUP98::HOXA9, or NUP98::LNP1 and HAT complex proteins (Fig 2C).

**Figure 2.**
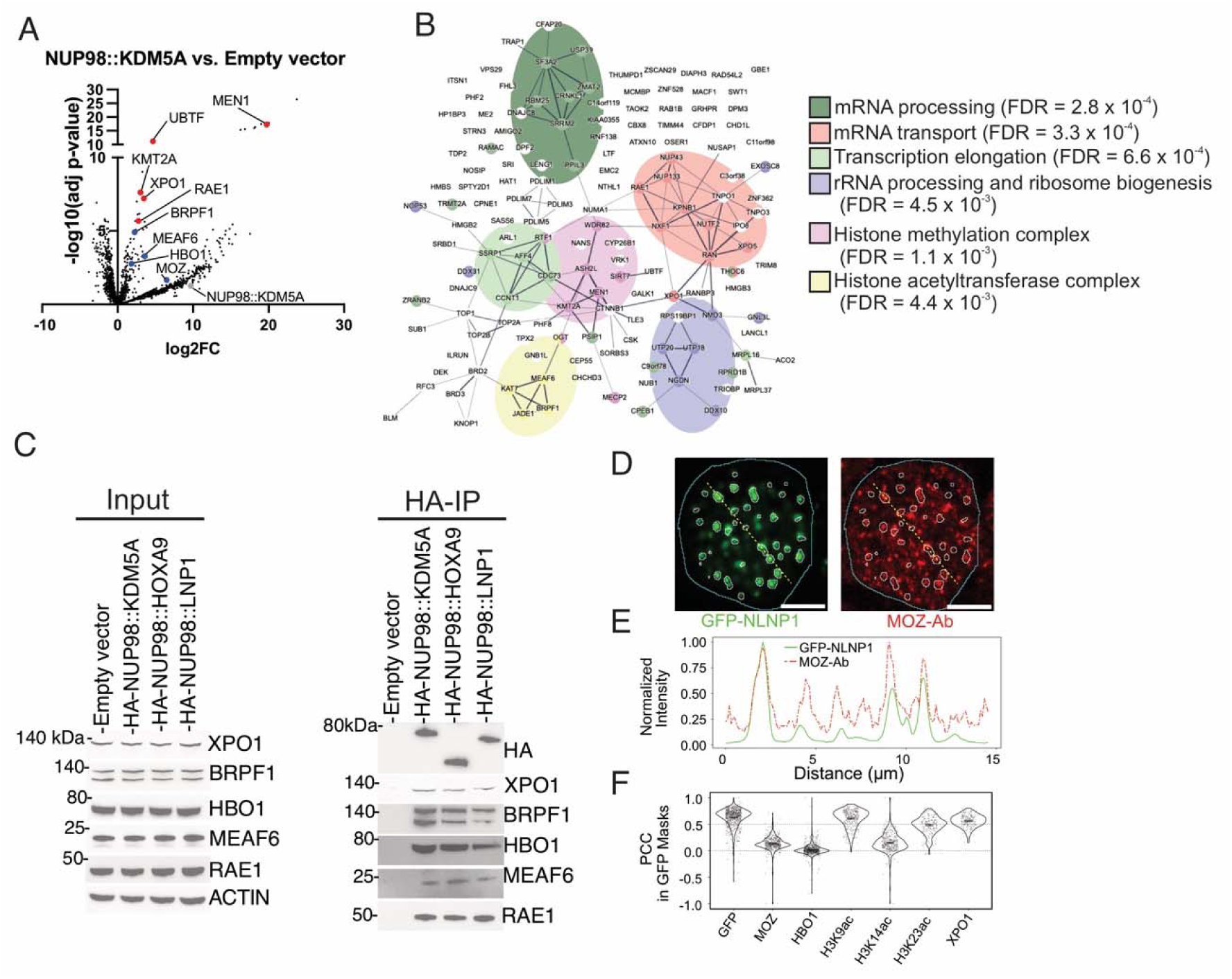
NUP98 FOs interact with histone acetyltransferase complex members. A) Volcano plot for enriched proteins in transiently transfected HEK293T cells expressing HA-NUP98::KDM5A compared to empty vector. The fusion bait protein and significantly enriched associated-proteins (adjusted *P*<0.05) are labeled. Data with missing values in the empty vector were imputed; these appear as a distinct cluster of high log_2_ fold-change values. The average log_2_ fold-change of all three biological replicates was used for the plot. B) STRING analysis of enriched proteins (log_2_ fold change > 1, adjusted *P*<0.025) from RIME in transiently transfected HEK293T cells expressing HA-NUP98::KDM5A (compared to empty vector). C) Western blot of input and IP samples for NUP98 FO interactors in transiently transfected HEK293T cells expressing empty vector or HA-NUP98 FO. D-F) HEK293T cells were transiently transfected with GFP-tagged NUP98::LNP1, fixed with methanol, and stained for HAT complex members or associated histone modifications. Images were acquired by confocal microscopy. D) Example cell showing antibody florescence signals for GFP-NUP98::LNP1 in green and MOZ in red. GFP condensate segmentation is shown with white outlines, and cell nuclear boundaries are shown with cyan outlines. The fluorescence intensities along the yellow lines were normalized and plotted in E. Scale bar, 5 μM. E) Line plot comparing normalized intensities of GFP and RRX signals along the dotted lines shown in D. F) Quantification of Pearson correlation coefficient (PCC) values for GFP-NUP98::LNP1 vs. Ab-RRX signals within GFP condensates. Values are shown for antibodies against the indicated proteins with each distinct point representing a single cell.

Based on previous work defining the importance of liquid-liquid phase separation for NUP98 FO-driven cell transformation and gene dysregulation (17,18), we hypothesized that the interacting proteins we identified by RIME may colocalize with NUP98 FOs in nuclear condensates. We transfected HEK293T cells with GFP-tagged NUP98::LNP1, which forms highly distinct condensates amenable to quantitative analysis. Our previous work showed that HEK293T cells form NUP98 FO condensates similar to those observed hematopoietic cells (18). After fixation, we stained cells with primary antibodies directed to putative NUP98 FO interactors. We used XPO1 antibody, as XPO1 was one of the top hits from RIME and is known to interact with NUP98 FOs (20,36,39), as well as GFP antibody as a positive control. We also stained cells with antibodies targeting MOZ, HBO1, and the acetylated histone marks deposited by these complexes, including H3K9ac, H3K14ac, and H3K23ac. Rhomadine Red-X (RRX)-conjugated secondary antibodies were used for visualization of these primary antibody targets. We examined the colocalization of antibody (red) signal with the FO (GFP) signal within FO condensates in 100+ cells for each condition by computing the Pearson correlation coefficient (PCC) for the red and green signals (Figure 2D-F, SFig 3F-K). We found that NUP98::LNP1 condensates showed high PCC values for FO and antibody signals for GFP, XPO1, H3K9ac, and H3K23ac antibodies (SFig 3F-G, I). We observed lower correlations with MOZ, HBO1, and H3K14ac signals (SFig 3H, J-K), but were still able to identify condensates in which colocalization did occur. Our results suggest that epigenetic regulators identified by RIME are present with NUP98 FOs in transcriptional condensates and may play an active role in leukemic gene regulation.

### MYST family HAT complex members are molecular dependencies in NUP98-r leukemia

To systemically determine epigenetic dependencies in *NUP98*-r leukemogenesis in vivo, we performed a CRISPR/Cas9 screen with a gRNA library targeting 338 genes involved in epigenetic modifications (40). Lin-HSPCs from Cas9 transgenic mice were transduced to express NUP98::KDM5A and gRNA, then were subsequently transplanted into lethally irradiated congenic recipient mice. After nine weeks, mice developed clinical signs of leukemia development. At the time of sacrifice, we isolated myeloid (leukemic) and T-cells (as an internal control) from the bone marrow and spleen. We performed deep sequencing to identify depleted gRNAs in tumor versus non-tumor cells, which nominated positive regulators required for NUP98::KDM5A leukemogenesis (Figure 3A, SFig 4A). One of the top hits from this in vivo screen was the epigenetic reader and MYST HAT complex member *Brpf1*, as the gRNAs targeting this gene were significantly depleted in both spleen and bone marrow samples (Figure 3B, SFig 4B, Table S7-S8). Furthermore, pathway analysis revealed that gRNAs targeting genes involved in the regulation of stem cell division and histone acetylation were positive regulators of NUP98::KDM5A FO-driven leukemia (Figure 3C, Table S9), consistent with our hypothesis that HAT complexes have a central role in *NUP98-*r leukemogenesis.

**Figure 3.**
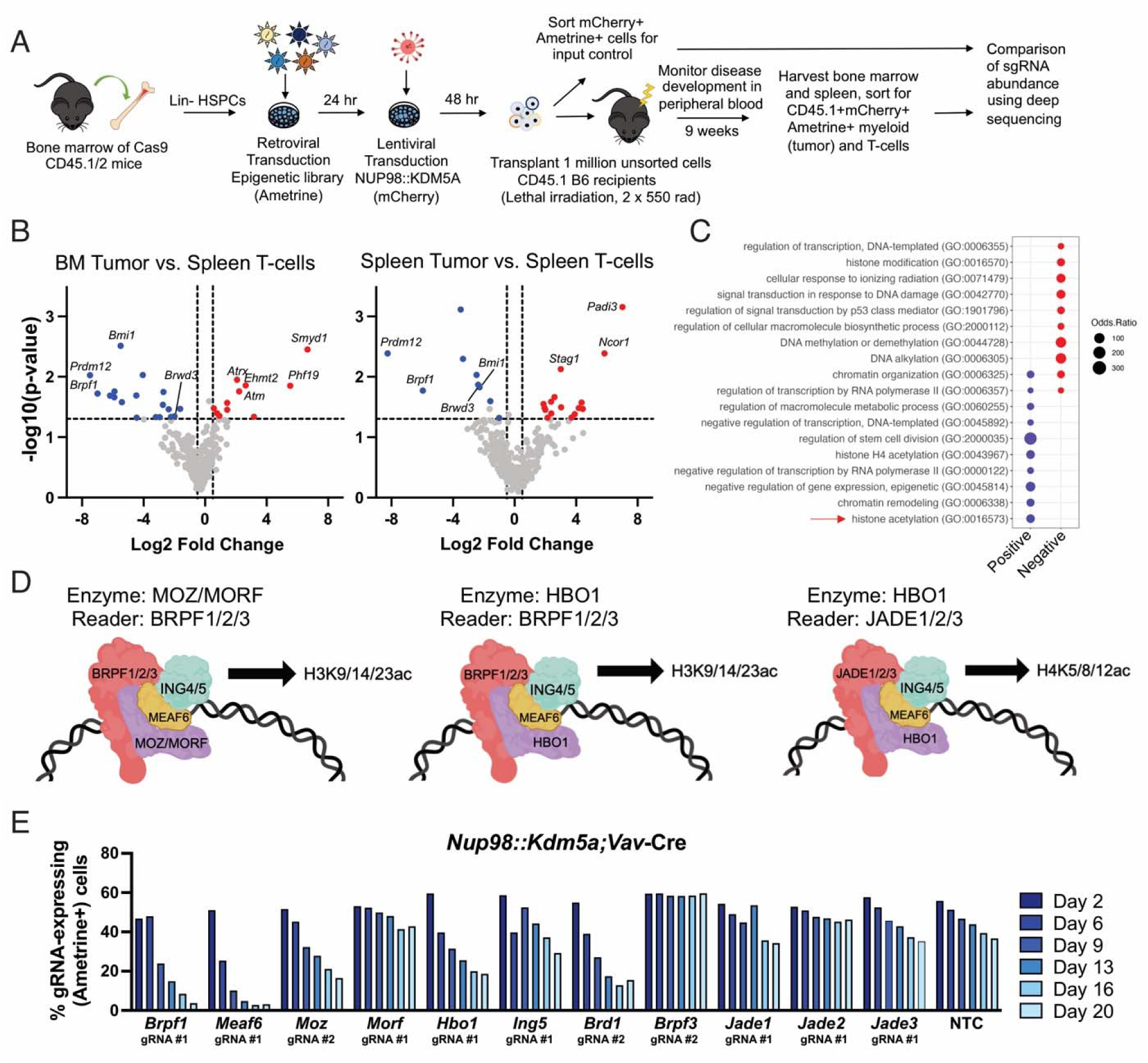
Histone acetyltransferase complex members are molecular dependencies in *NUP98*-r cells. A) Schematic of in vivo epigenetic CRISPR screen. B) Volcano plot for enriched/depleted gRNAs in bone marrow and spleen cells of NUP98:KDM5A mice vs. spleen T-cell controls from the same animals. C) Gene set enrichment analysis of enriched/depleted gRNAs. D) Schematic of BRPF1-containing histone acetyltransferase complexes and the histone H3 or H4 acetylation modifications that each mediates. E) Relative fitness of *Nup98::Kdm5a;Vav*-Cre;Cas9 HSPCs transduced with gRNAs targeting HAT complex members (marked with Ametrine) and placed in competitive with similar cells transduced with non-targeting control gRNA (marked with GFP). Representative data is shown, and a second replicate is in SFig 4C.

To validate these results, we performed CRISPR competition assays using NUP98 FO-expressing Cas9+ lin-HSPCs, which were transduced to express gRNAs targeting a gene of interest (Ametrine+) and placed in competition with control gRNA-expressing (GFP+) in culture. We used this strategy to assess how genetic disruption of MYST family HAT complex members (Figure 3D) affects *NUP98*-r cell fitness (41). *Nup98::Kdm5a;Vav-*Cre;Cas9 lin-HSPCs expressing gRNAs targeting *Brpf1* were rapidly and profoundly depleted, consistent with the results of our in vivo CRISPR screen (Figure 3E, SFig 4C). We validated this finding in Cas9+ retroviral models of NUP98::LNP1 (SFig 4D). Moreover, gRNAs targeting other members of BRPF1-containing HAT complexes (Figure 3D), such as *Meaf6, Moz*, *Hbo1*, or *Brd1*, also led to decreased cell fitness. Our results point to a specific molecular dependency on MOZ- and HBO1-containing HAT complexes, since genes involved in other HAT complexes (*Morf, Ing5*, *Jade1/2/3, Brpf3*) did not impact the proliferation of *NUP98*-r cells (Figure 3E, SFig 4D). These results prompted us to focus on H3 acetylation changes, as BRPF readers are specifically associated with H3 histone acetylation, whereas JADE proteins read H4 modifications (42). Altogether, our in vivo screen and CRISPR competition assay validation suggest that MYST family HAT complexes containing MOZ and/or HBO1 have critical roles in sustaining *NUP98*-r leukemogenesis, potentially through the regulation of H3 acetyl modifications that promote an active chromatin landscape.

### Chemical HAT complex inhibition leads to loss of acetylated histone modifications and myeloid differentiation in NUP98-r leukemia

MOZ and HBO1 have been associated with acetylation of histone H3 at lysine 9, 14, and 23 in prior literature (43–48). Nevertheless, the biochemical roles of MOZ, MORF, and HBO1 on individual histone acetylation modifications remains to be fully determined, and most lysine acetyltransferase inhibitors have modest specificity for MOZ over other MYST family HATs. Recent data highlight the contribution of MYST family HATs in H3K14 and H3K23 acetylation (44–48), even suggesting that these acetyl modifications are changed on the same histone tails after MORF disruption (47). However, there are few antibodies with demonstrated specificity for H3K23ac, and those commercially available are not suitable for chromatin immunoprecipitation or CUT&RUN studies, particularly in mouse cells (data not shown). These technical limitations hinder mechanistic understanding of the specific role of epigenetic changes in *NUP98*-r pathobiology.

To overcome these limitations and systematically evaluate the impact of MYST HAT inhibitors on histone posttranslational modifications (PTMs), we performed mass spectrometry in *NUP98*-r cells treated with MOZ/HBO1 small molecule inhibitor PF9363. PF9363 is a potent, orally bioavailable inhibitor of MOZ/HBO1 histone acetyltransferase activity, which is being evaluated in early phase clinical trials (46,49). PF9363 inhibits MOZ activity at low nanomolar concentrations and HBO1 at higher nanomolar concentrations (46). We isolated histones from *Nup98::Kdm5a;Vav-*Cre;Cas9 HSPCs treated for 72 hours with DMSO or PF9363 and performed mass spectrometry to identify histone PTMs, or combinations of PTMs, that were significantly increased or decreased between treatment groups. At the 1 μM dose of PF9363, targeting both MOZ and HBO1, we found that H3K23ac was the most significantly reduced histone PTM, with H3K14ac alone or in combination with H3K9me1, H3K9me2, or H3K9me3 being reduced to a lesser extent (Figure 4A). Titrating the dose of PF9363 down to 10 nM, now targeting only MOZ, maintained the decrease in H3K23ac, while H3K14ac was no longer changed with lower concentrations of PF9363 (Figure 4B, SFig 5A). In contrast, the overall abundance of H3K9ac was not significantly changed even after 1 μM PF9363 treatment (Figure 4H). Overall, these results are consistent with previous observations that MOZ disruption leads to locus-specific reduction of H3K9ac (43) and global reduction of H3K23ac (47). Furthermore, the more profound loss of H3K23ac is in agreement with work by Sharma *et al.*, which suggests that H3K23ac may be the primary target of MOZ and that H3K14ac is mediated by HBO1 and would only be targeted at higher concentrations of PF9363 (46).

**Figure 4.**
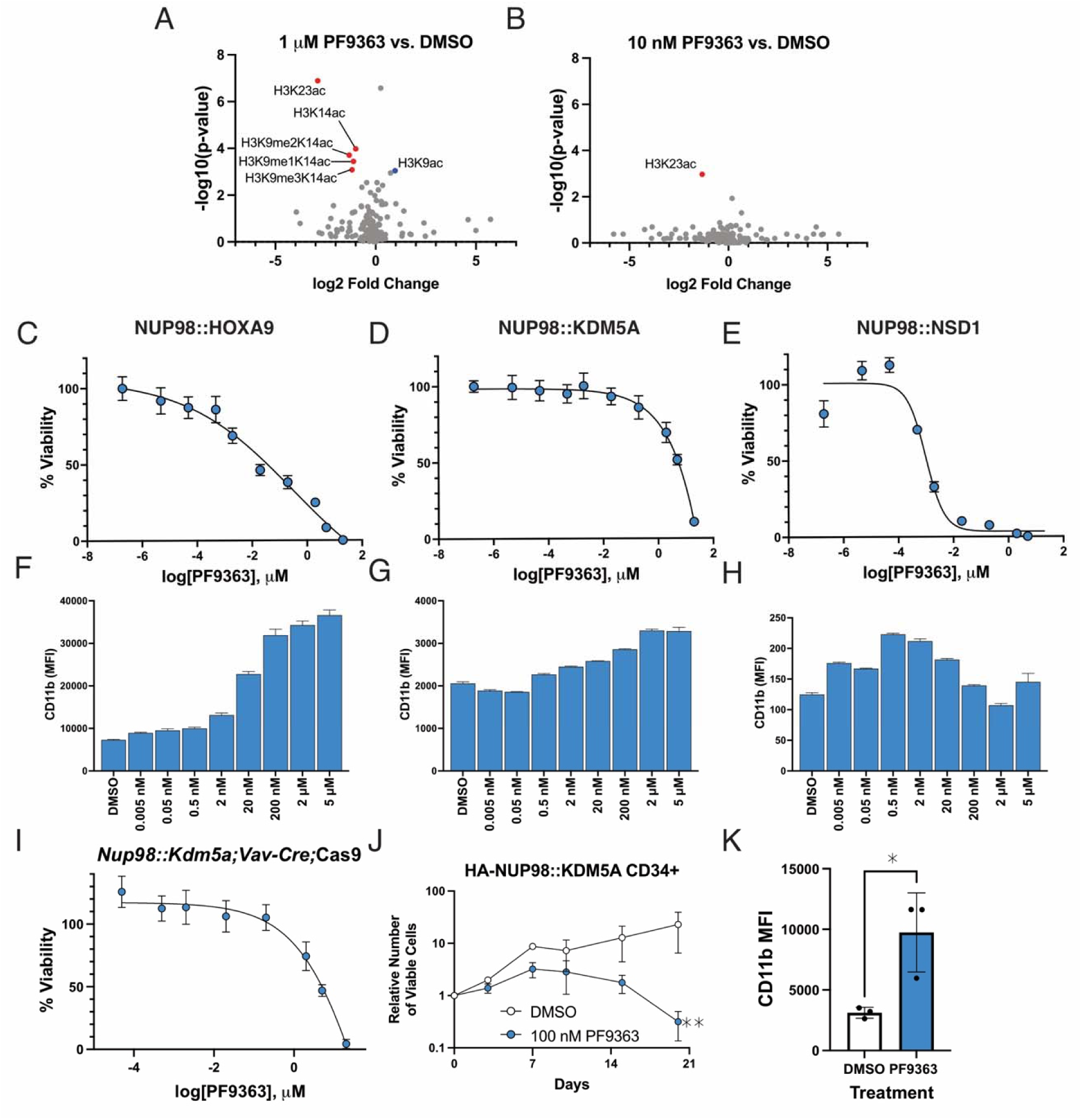
Pharmacologic inhibition of MOZ/HBO1 with PF9363 results in loss of acetylated histone H3 modifications and negatively impacts *NUP98*-r cell fitness. Volcano plots of histone PTMs identified by mass spectrometry after treatment of *Nup98::Kdm5a;Vav*-Cre-Cas9 HSPCs for 72 hours with vehicle (DMSO) or A) 1 μM or B) 10 nM PF9363. C) NUP98::HOXA9, D) NUP98::KDM5A, and E) NUP98::NSD1 mouse leukemia cells were treated with MOZ inhibitor PF9363 for 10 days, and viability was measured compared to vehicle (DMSO)-treated controls. F) NUP98::HOXA9, G) NUP98::KDM5A, and H) NUP98::NSD1 mouse leukemia cells were treated with MOZ inhibitor PF9363 for 10 days, and CD11b expression was measured compared to vehicle (DMSO)-treated controls. I) *Nup98::Kdm5a;Vav*-Cre;Cas9 HSPCs were treated with increasing concentrations of PF9363 for 9 days and viability was measured using a resazurin cell viability assay. CD34+ HSPCs transduced to express HA-NUP98::KDM5A were cultured in media containing vehicle (DMSO) or 100 nM PF9363 for 19 days. J) Growth curve with mean +/- SD and mixed linear model, ** denotes *P*<0.01. K) CD11b expression measured by flow cytometry at day 19. Mean +/- SD and unpaired t-test, * denotes *P*>0.05. Experiments were performed in technical triplicate, representative data from one of two biological replicates (different CD34+ cell donors) is shown.

We hypothesized that PF9363-induced loss of acetylated histone modifications would impact key pro-leukemogenic genes, thus leading to decreased leukemia cell fitness; we therefore evaluated the therapeutic impact of pharmacologic inhibition of MOZ and HBO1 in *NUP98*-r cells. Consistent with what has been previously reported for other small molecules targeting epigenetic regulators (50,51), we found that extended treatment (10 days) with PF9363 inhibited the growth of multiple *NUP98*-r models, including cells expressing NUP98::HOXA9,::KDM5A, or::NSD1 (Figure 4C-E). Indeed, treatment with PF9363 demonstrated similar efficacy to the Menin inhibitor revumenib, which is being evaluated in clinical trials of *NUP98*-r AML (NCT04065399) (24,52). Moreover, treatment with PF9363 was accompanied by a modest increase in CD11b expression (Figure 4F-H), indicative of myeloid differentiation. HSPCs from *Nup98::Kdm5a;Vav-*Cre mice were also responsive to 10-day treatment with PF9363 (Figure 4I).

To evaluate the effects of MOZ/HBO1 inhibition in human models of *NUP98* rearrangement, we transduced cord blood CD34+ cells to express HA-NUP98::KDM5A and treated them with vehicle (DMSO) or 100 nM PF9363 for 3 weeks. HA-NUP98::KDM5A CD34+ cells were also responsive to PF9363, as treatment led to reduced cell growth (Figure 4J).

Moreover, treatment with 100 nM PF9363 led to increased expression of CD11b (Figure 4K), similar to our observations in the NUP98 FO mouse cells (Figure 4F-H).

We also investigated the effects of PF9363 on normal hematopoietic cells. We treated human peripheral blood mononuclear cells (PBMCs) with PF9363 for 72 hours, and we observed limited reduction in cell viability only at high concentrations (>1 μM) (SFig 5B). In CFU assays with human cord blood CD34+ cells, PF9363 treatment led to a 20-30% reduction in the total number of colonies, but all colony lineages were maintained. Together, these data suggest that there is a therapeutic window for MOZ/HBO1 inhibitors in hematopoietic cells.

### MOZ inhibition decreases leukemic burden in NUP98-rearranged PDXs

Given the strong dependency of *NUP98-*r leukemia cells on MYST HAT activity, we hypothesized that pharmacologic inhibition of MOZ/HBO1 would be an effective therapeutic strategy for *NUP98*-r patient-derived xenograft (PDX) models in vivo. Mice bearing a PDX model of chronic myeloid leukemia bearing *NUP98::HOXA13* and *BCR::ABL1* (SJCML068699) (53) received vehicle or 75 mg/kg SNDX-5613 twice daily, or 3 mg/kg PF9363 daily for 5 days each week for 6 weeks. Treatment with PF9363 led to decreased leukemic burden, and this effect was sustained even after the end of the 6-week treatment period (Fig 5A, SFig 6A-C). Notably, this PDX was completely resistant to SNDX-5613 treatment, suggesting that MOZ/HBO1 HAT inhibitors may have non-overlapping targets with KMT2A-Menin inhibitors in some *NUP98*-r leukemias (SFig 6D).

**Figure 5.**
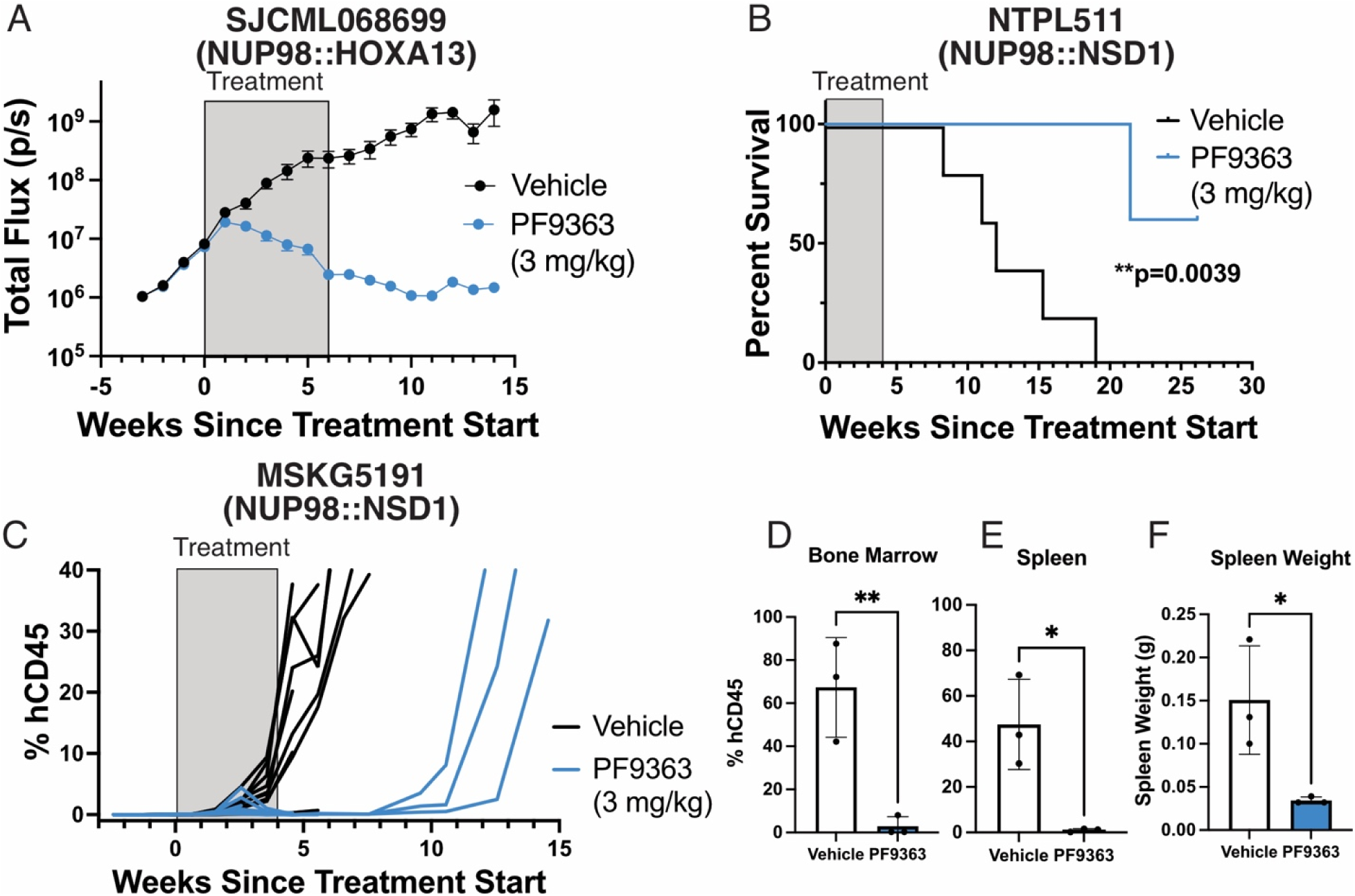
*NUP98*-r PDXs respond to MOZ inhibition in vivo. A) Total ventral flux was monitored over time for mice bearing luciferase-marked SJCML068699 (NUP98::HOXA13) PDX (n=10/group) and treated with vehicle or PF9363 (3 mg/kg, 5 days on, 2 days off, IP) for 6 weeks. B) Survival curve for mice bearing NTPL511 (NUP98::NSD1) PDX (n=5/group) treated with vehicle or PF9363 (3 mg/kg, QD, IP) for 4 weeks. For mice bearing MSKG5191 (NUP98::NSD1) PDX, human CD45 was monitored over time in the C) peripheral blood and at the end of 4-week treatment in the D) bone marrow and E) spleen. F) Spleen weight for MSKG5191 PDX-bearing mice at the end of treatment. For D-F, mean±SD are shown, and using unpaired t-test, * denotes *P*>0.05, ** denotes *P*<0.01.

We expanded these findings in two additional *NUP98*-r PDX models harboring the most common translocation partner seen in patients: NUP98::NSD1 (NTPL511 and MSKG5191). In the NTPL511 model, which we had previously shown is completely responsive to Menin inhibitor revumenib (24), mice were treated daily for 4 weeks with vehicle or 3 mg/kg PF9363 and then monitored to determine if treatment would lead to a survival benefit. All vehicle-treated mice developed leukemia and succumbed to disease before any PF9363-treated mice became sick (Figure 5B, *P*=0.0039). Nevertheless, in two of five (40%) PF9363-treated mice, leukemia eventually reemerged, and at the time of sacrifice, vehicle- and PF9363-treated animals displayed no difference in the percentage of human leukemia cells in the blood or bone marrow and no difference in spleen weight (SFig 6E-G). A second PDX, MSKG5191 was derived from a heavily pretreated pediatric AML patient harboring NUP98::NSD1 as well as *FLT3-*ITD, *BCORL1, ASXL1*, *IDH1* and *WT1* mutations, and was unresponsive to Menin inhibition with revumenib (24). Four weeks of daily treatment with 3 mg/kg PF9363 significantly reduced the leukemia burden in the blood, bone marrow and spleen, although just as in the NTPL511 model, leukemia reemerged after cessation of PF9363 treatment (Figure 5C-F). Taken together, these results indicate that MOZ inhibition is a promising therapeutic strategy in *NUP98*-r leukemia, but these results also suggest that combination therapies may be required to fully eradicate the disease.

### Efficacy of combined MOZ/HBO1 and Menin inhibition

Based on these findings and the reemergence of leukemia after MOZ/HBO1 inhibitor treatment in two of three PDXs that we tested, we investigated whether MOZ/HBO1 and Menin inhibitor combination therapy would be more effective than either agent alone. We treated *Nup98::Kdm5a;Vav*-Cre HSPCs with MOZ inhibitor PF9363, Menin inhibitor SNDX-5613 (revumenib), or combination for 10 days and observed a synergistic effect when both drugs were used together with a Bliss synergy score of 44.5 (Figure 6A-B, SFig 7A). We also observed that the combination of 1 μM PF9363 and 1 μM SNDX-5613 inhibited self-renewal of

**Figure 6.**
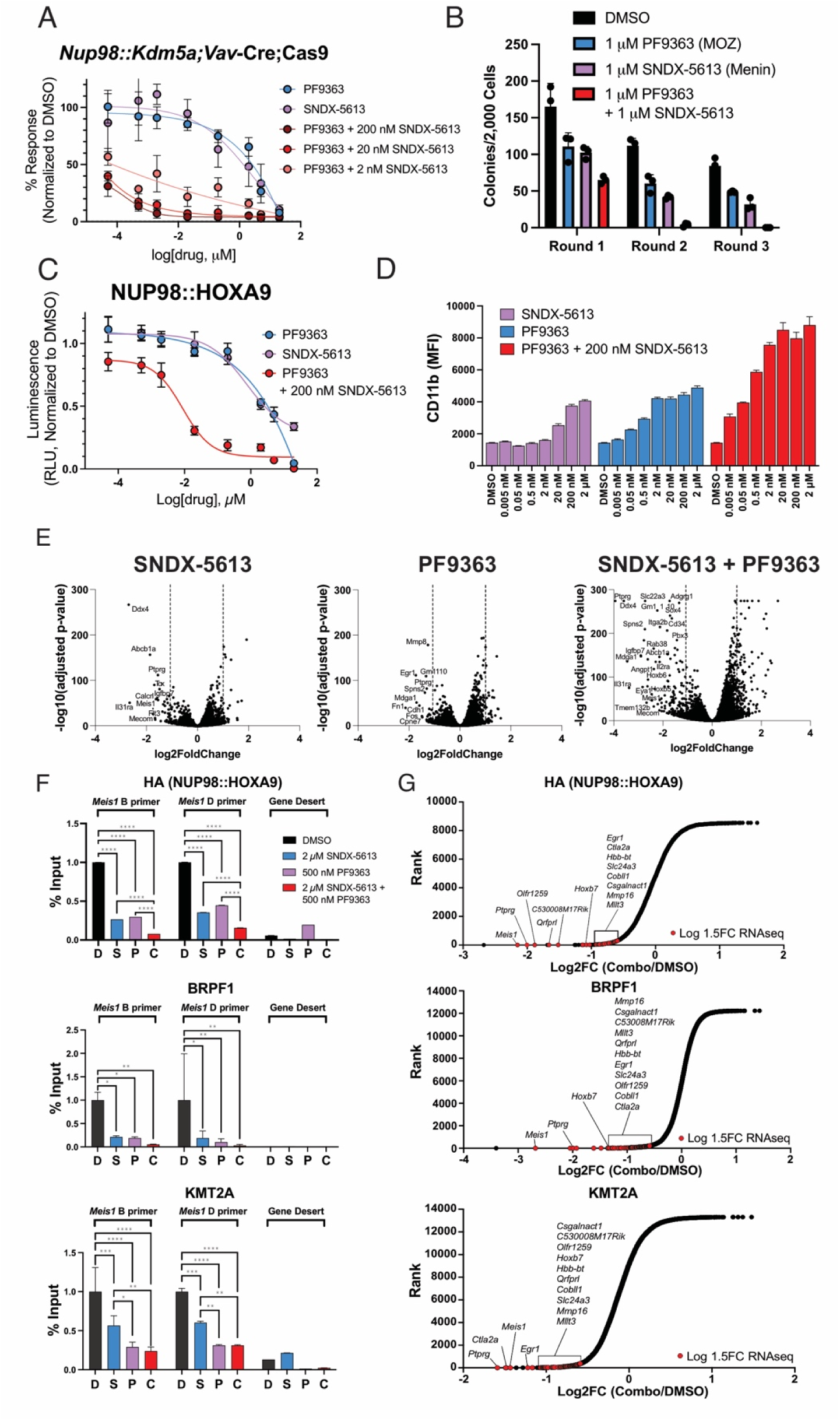
MOZ/HBO1 and Menin inhibition alter gene expression and remodel chromatin in *NUP98*-r cells. A) *Nup98::Kdm5a;Vav*-Cre;Cas9 HSPCs were treated with PF9363, SNDX-5613, or combination for 9 days, and viability was measured relative to vehicle (DMSO)-treated controls by flow cytometry for DAPI. B) Colony forming unit assay for *Nup98::Kdm5a;Vav*-Cre lin-HSPCs treated with DMSO, 1 μM PF9363, 1 μM SNDX-5613, or combination. NUP98::HOXA9 mouse leukemia cells were treated with PF9363, SNDX-5613, or combination for 10 days, and C) viability (using DAPI staining) or D) CD11b expression was measured by flow cytometry relative to vehicle (DMSO)-treated controls. E) Volcano plot showing differentially expressed genes in RNA-seq samples from NUP98::HOXA9 mouse leukemia cells treated for 72 hours with 2 μM SNDX-5613, 500 nM PF9363, or combination relative to vehicle (DMSO)-treated cells. F) ChIP-qPCR at the *Meis1* locus for HA (NUP98::HOXA9), KMT2A, and BRPF1 in NUP98::HOXA9 mouse leukemia cells after 72-hour treatment with DMSO, 2 μM SNDX-5613, 500 nM PF9363, or combination. Mean +/- SD by 2-way ANOVA and Tukey’s multiple comparison test, * denotes *P>*0.05, ** denotes *P*<0.01, *** denotes *P*<0.001, and **** denotes *P*<0.0001. G) Hockeystick plots depict log_2_ fold ratio of chromatin occupancy for each ChIP target after 72-hour treatment with 2 μM SNDX-5613 and 500 nM PF9363 with gene rank ordered by chromatin occupancy. Targets with gene expression changes greater than 1.5 log_2_ fold change reduction after treatment are highlighted in red. Common targets in HA, BRFP1, and KMT2A ChIP experiments are annotated. Representative data from n = 3 independent experiments is shown.

*Nup98::Kdm5a;Vav-Cre* HSPCs in a CFU assay (Figure 6B). Cells treated with SNDX-5613 or PF9363 inhibitor alone formed fewer colonies than cells treated with vehicle, and immunophenotyping of SNDX-5613- or PF9363-treated colonies revealed increased expression of myeloid differentiation markers CD11b and Gr1 (SFig 7B). Similar results were observed with the MOZ/HBO1 inhibitor WM-1119 and the Menin inhibitor VTP-50469 (revumenib) (SFig 7C-D). To validate these findings in NUP98 FO leukemias with different fusion partners, we treated NUP98::HOXA9 and NUP98::NSD1 FO-expressing mouse leukemia cells with PF9363, SNDX-5613 and combination. Combined treatment was synergistic with a Bliss synergy score of 24 and increased CD11b expression beyond monotherapies (Fig 6C-D, SFig 7E-F).

To understand the molecular mechanisms underlying response to MOZ/HBO1 and Menin inhibition in *NUP98-*r cells, we performed RNA-seq and ChIP on NUP98::HOXA9 mouse leukemia cells after 72-hour treatment with DMSO, 500 nM PF9363, 2 μM SNDX-5613, or combination. While some genes were downregulated after monotherapy SNDX-5613 or PF9363, we observed the most robust changes in gene expression with combination treatment, including significantly decreased expression of NUP98 FO target genes, including *Meis1* and the *Hoxb* gene cluster (Figure 6E, SFig 7G). GSEA revealed that combination treatment led to the downregulation of stem cell-associated genes and upregulation of monocyte and granulocyte-monocyte progenitor (GMP) gene sets (SFig 7H, Table S10, Table S11). These results suggest that Menin-KMT2A and MOZ/HBO1 may cooperatively regulate a subset of NUP98 FO target genes, whose expression is further attenuated by combination therapy.

Consistent with these results, we observe that NUP98 FO, KMT2A, and BRPF1 co-localize near the transcriptional start site (TSS) of key pro-leukemogenic genes like *Meis1* (SFig 7I). We sought to further define how treatment with small molecule inhibitors of Menin and MOZ/HBO1 impacted chromatin occupancy of these complexes. Our previous work has demonstrated that pharmacologic targeting of Menin in *NUP98-*r leukemia cells results in chromatin changes at transcriptionally sensitive loci. Similar to what we have reported for Menin inhibitors, treatment of *NUP98*-r leukemia cells with PF9363 did not produce changes in global chromatin occupancy of the NUP98 FO or the common MOZ/HBO1 complex component BRPF1 (SFig 7J). Rather, treatment with PF9363 resulted in loss of the NUP98 FO, BRPF1, and KMT2A at transcriptionally sensitive genes that were downregulated in our RNA-seq studies. For example, treatment with 500 nM PF9363 resulted in loss of chromatin occupancy of NUP98 FO (marked with HA epitope tag), BRPF1, and KMT2A (in complex with Menin) at the *Meis1* locus (Figure 6F, SFig 7I). Consistent with what we have previously published, we observed loss of the NUP98 FO and KMT2A chromatin occupancy upon treatment of *NUP98*-r leukemia cells with SNDX-5613 (Figure 6F, SFig 7I). By contrast, genes such as those in the *Hoxa* cluster, which do not change expression upon treatment with SNDX-5613 or PF9363 (Figure 6E), demonstrated none of these chromatin changes (Figure 6F, SFig 7K). Strikingly, combination treatment with PF9363 and SNDX-5613 resulted in more profound loss of the NUP98 FO chromatin occupancy at certain NUP98 FO target genes, such as *Meis1* and *Hoxb7* (Figure 6G). Altogether, these results demonstrate that MOZ/HBO1 and Menin regulate a partially overlapping but also distinct subset of NUP98 FO pro-leukemogenic target genes. Combined pharmacologic inhibition of these complexes, therefore, may synergistically impact gene expression at shared target loci and many function to promote differentiation.

### Combined MOZ and Menin inhibition overcomes resistance to Menin inhibition

Since our RNA-seq and ChIP-seq studies revealed that combined MOZ/HBO1 and Menin inhibition could cooperatively displace NUP98 FO from chromatin, we hypothesized that combination treatment could be effective in *NUP98-*r PDX models in which Menin inhibitors have limited therapeutic efficacy. For example, we have previously reported that a NUP98::KDM5A acute megakaryocytic leukemia PDX model (CPCT0021) responds to Menin inhibitor therapy, but mice subsequently relapse weeks after withdrawal of drug. We treated mice engrafted with PDX cells from this model with vehicle, PF9363 (IP, QD), SNDX-5613 chow, or a combination of both agents (n = 5/group), and evaluated disease burden in each treatment group at the time of clinically manifest disease in control mice. Treatment with SNDX-5613, PF9363 or combination led to a significant decrease in the percentage of human CD45+ cells in the spleen as well as a reduction in spleen weight (Figure 7A, SFig 8A). Disease burden was also reduced for each treatment in the peripheral blood, although only SNDX-5613 led to a reduction in human CD45+ cells in the bone marrow (Figure 7A). We evaluated CD41 and CD61 expression as indicators of megakaryocytic cell differentiation, and we observed that combination treatment resulted in the most robust upregulation of these mature myeloid markers in the bone marrow and peripheral blood (Figure 7B, SFig 8B). To further consider how MOZ inhibition affects gene expression in this PDX, we performed RNA-seq on samples collected at the end of the treatment period from vehicle- and PF9363-treated animals. We observed upregulation of megakaryocytic lineage genes including von Willebrand factor (*VWF*) and multiple integrins (*ITGB3*, *ITGA2*, *ITGB5*) (Figure 7C). GSEA revealed upregulation of genes with increased expression after Menin inhibitor treatment of the same PDX model (24) and of genes involved in megakaryocytic differentiation (Figure 7D, Table S12). Downregulated genes and gene sets include those with decreased expression after Menin inhibition (24) and those expressed in myeloid cells, again providing evidence of megakaryocytic differentiation (SFig 8C-D, Table S13). Immunohistochemistry on bone marrow samples from treated mice showed increased expression of human CD45 in PF9363-, SNDX-5613, and combination-treated animals. These results suggests that treated PDX cells are more mature than those from vehicle-treated mice, which exhibit low expression of human CD45 and a more primitive phenotype (SFig 8E). Staining for VWF by immunohistochemistry (IHC) was increased in drug-treated animals, consistent with the upregulation of *VWF* in RNA-seq data (Figure 7C, SFig 8E). Next, we asked whether the addition of MOZ/HBO1 inhibitor could overcome Menin inhibitor resistance in a NUP98::NSD1 PDX (MSKG5191) that is completely unresponsive to Menin inhibitor VTP-50469 (24). Following engraftment of PDX cells, we treated animals for 4 weeks with vehicles, 3 mg/kg PF9363 (IP, daily), 75 mg/kg SNDX-5613 (PO, twice daily), or combination (n = 10/group). We sacrificed half of the mice at the end of treatment, which revealed decreased spleen weight in the PF9363- and combination-treated arms (SFig 8E). Further, the combination-treated mice had lower disease burden in the bone marrow and spleen as well as a significant increase in the percentage of CD11b-expressing (differentiated myeloid) cells in both tissues (SFig 8F). Increased staining for human CD45 and CD33 was observed by IHC in the sternum, which also indicates that combination-treated cells undergo myeloid differentiation (SFig 8G). Interestingly, the hCD45 population in the peripheral blood decreased dramatically only after the end of treatment (Figure 7E), which is likely a result of the differentiation noted by immunophenotyping. Half of the mice were subject to disease monitoring after the end of 4-week treatment, to determine if treatment with PF9363 and/or SNDX-5613 would lead to a survival benefit. We observed that PF9363 led to a 21% increase in disease latency compared to vehicle (Figure 7F). Moreover, combination-treated mice survived significantly longer than mice from all other treatment groups (Figure 7F). Although all mice did eventually succumb to leukemia, combined treatment with PF9363 and SNDX-5613 resulted in a 50% increase in disease latency compared to vehicle-treated animals and 24% increase compared to PF9363 monotherapy treatment.

**Figure 7.**
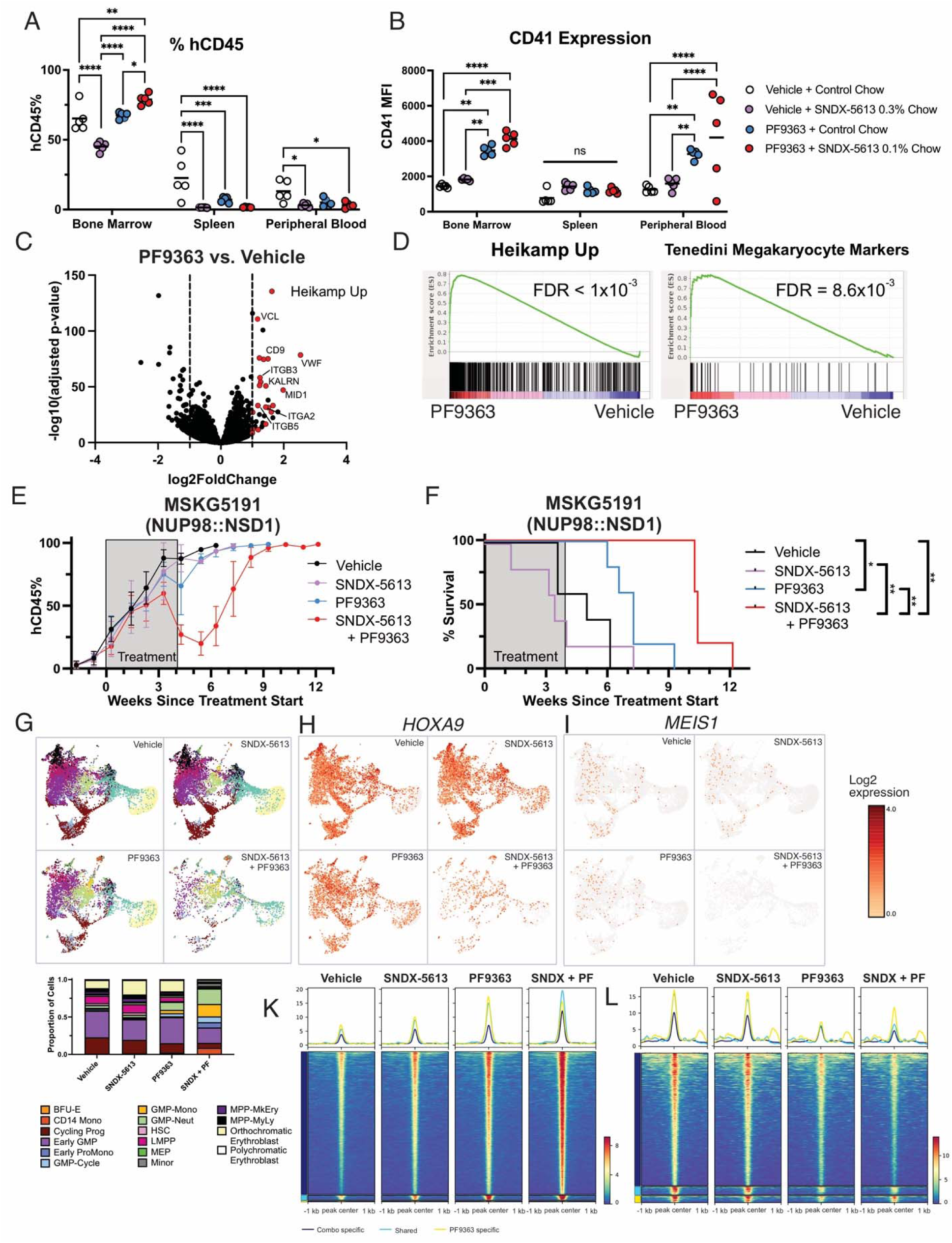
MOZ/HBO1 and Menin inhibition have combinatorial effects in vivo. Mice bearing CPCT0021 (NUP98::KDM5A) PDX cells were treated for 30 days with vehicles, PF9363 (3 mg/kg, QD, IP), SNDX-5613 (chow), or combination. A) Human CD45 or B) CD41 expression was measured at the end of treatment in the bone marrow, spleen, and peripheral blood. Statistical comparisons were performed using 2-day ANOVA. C) Volcano plot showing differentially expressed genes in RNA-seq samples from vehicle or PF9363-treated mice. D) Gene set enrichment analysis showing upregulation of Heikamp Up and Tenedini Megakaryocyte Markers gene sets in NUP98::KDM5A PDX cells after PF9363 treatment. Mice bearing NUP98::NSD1 PDX cells were treated for 4 weeks (5 days on, 2 days off) with vehicles, PF9363 (3 mg/kg, QD, IP), SNDX-5613 (75 mg/kg, BID, PO), or combination. E) Human CD45 was monitored over time in the peripheral blood. F) Survival curve for NUP98::NSD1 PDX-bearing mice during and after treatment. Statistical comparison was performed using Mantel-Cox test. Single cell RNA-seq was performed using the 10X Genomics 5’ technology in spleen samples from n = 2 mice per treatment arm (pooled samples). G) UMAP projection with clustering based on cell type and bar plot showing proportion of cells from each treatment group in each cluster. Cells that were not assigned to a cluster were omitted from this analysis/plot. BFU-E, burst forming unit-erythroid; Mono, monocyte; Prog, progenitors; GMP, granulocyte-monocyte progenitor; ProMono, promonocyte; Neut, neutrophil; HSC, hematopoietic stem cell; LMPP, lymphoid-primed multipotent progenitor; MEP, megakaryocyte-erythroid progenitor; MPP, multipotent progenitor; MkEry, megakaryocyte erythroid; MyLy, myeloid lymphoid. UMAP plots colored according to I) *HOXA9* and J) *MEIS1* expression between treatment arms. ATAC-seq was performed on human leukemia cells isolated from the spleens of MSKG5191 PDX-bearing mice after 5-day treatment with vehicle, SNDX-5613, PF9363, or combination. Tornado plots showing regions with K) differentially increased and L) differentially decreased accessibility between vehicle and treated samples. For all panels, * denotes *P*> 0.05, ** denotes *P*<0.01, *** denotes *P*<0.001, and **** denotes *P*<0.0001.

We hypothesized that combination treatment with MOZ/HBO1 and Menin inhibitors induced *NUP98-*r differentiation commensurate with the cell of origin for each PDX model. Therefore, we sought to investigate how cellular hierarchies were altered by MOZ and/or Menin inhibition in vivo by performing single cell RNA-seq (scRNA-seq). scRNA-seq from samples isolated after 4-week treatment showed distinctive changes between PDX cells from vehicle- and PF9363-treated mice, but the most striking changes in lineage trajectories were seen in cells from combination-treated animals. In leukemias treated with PF9363 monotherapy or combination, we observed an increase in the proportion of CD14 monocytes as well as all types of granulocyte-monocyte progenitors (GMPs) except early GMPs (Figure 7G, SFig 8I). These changes were accompanied by a decrease in hematopoietic stem cells and cycling progenitors (Figure 7G) and were consistent with the increase in myeloid differentiation noted by immunophenotyping and IHC above (Figure 7E, SFig 8J-K). Furthermore, we compared the expression of *NUP98* fusion target genes *HOXA9* and *MEIS1* between the four treatment groups. Treatment with PF9363 and SNDX-5613 led to a striking reduction in the expression of both target genes compared to the expression levels seen in vehicle or monotherapy treatments (Figure 7H-I, SFig 8L-M).

We then sought to understand how changes in the chromatin landscape were contributing to changes in gene expression and leukemic burden. To do so, we treated mice bearing MSKG5191 PDX cells with vehicle, SNDX-5613, PF9363 and/or combination treatment for 5 days and harvested leukemic spleen cells for assay for transposase-accessible chromatin (ATAC)-seq. Treatment with PF9363 led to modest changes in chromatin accessibility, while the addition of SNDX-5613 further enhanced the accessibility of many of the same loci with modest changes after PF9363 treatment (Figure 7J). Using motif analysis of differentially accessible regions, we identified that ETS factor motifs (i.e. SPI1, CEBPA, ETV6, ELF4) were overrepresented in regions with increased accessibility after treatment with SNDX-5613 and PF9363 (Table S14). Many of these transcription factors are also associated with myeloid differentiation, supporting the results of earlier experiments in this study. Conversely, regions with modest decreases in accessibility after PF9363 were less accessible in combination-treated cells (Figure 7K). These regions were enriched in GATA2 and RUNX1 motifs, which are associated with stem cell development and maintenance (Table S15). Overall, these scRNA-seq and ATAC-seq studies demonstrate that combined MOZ/HBO1 and Menin inhibitor treatment can reprogram stem cell-associated gene expression programs and drive leukemia cells to differentiate. Altogether, these studies provide mechanistic insight and preclinical justification for the synergistic combination of MOZ/HBO1 and Menin inhibitors in *NUP98*-r leukemia.

## DISCUSSION

Our results show that MYST family HAT complex members interact with NUP98 FOs in condensates and validate MOZ/HBO1 inhibition as an efficacious therapeutic strategy in *NUP98-*r leukemias. Using RIME to identify FO interactions that occur on chromatin, our studies identified interactions between HAT complex members and diverse NUP98 FOs (Figure 2C), suggesting that the NUP98 FO-HAT complex interaction involves the shared N-terminal portion of the FO. We also report colocalization of NUP98 FO with MOZ/HBO1 and acetylated histone H3 modifications within nuclear condensates, suggesting that the biochemical basis of these protein-protein interactions occurs in the context of phase-separated condensates. The variability in colocalization of HAT complex members with NUP98 FO suggest that these interactions may be weak or transient, or it may indicate that MOZ/HBO1 is present within a subset of condensates, which may have a unique contribution to *NUP98*-r phenotypes. Future work will aim to dissect whether pharmacologic inhibition of MOZ/HBO1 and loss of H3 acetyl modifications alters condensate composition or function, and whether disruption of FO condensates contributes to the therapeutic effect of these small molecule inhibitors.

Our studies were enabled by the generation and use of conditional knock in mouse models of *Nup98::Kdm5a* and *Nup98::Nsd1*. Previous work has investigated transgenic models of NUP98::HOXD13 as well as multiple other NUP98 FOs (25,54–57), but to our knowledge, genetically engineered mouse models of NUP98::KDM5A and NUP98::NSD1, with fusion gene expression driven by the endogenous *Nup98* promoter at the physiologic level rather than by ectopic overexpression, have not been described. Previous studies, including those from our group (14,58), used viral overexpression of NUP98::KDM5A to develop in vivo leukemia, and our data from *Nup98::Kdm5a;Vav-*Cre mice largely recapitulates these findings. Although our *Nup98::Kdm5a* conditional knock in model leads to acute leukemia with latency comparable to previous genetically engineered mouse models of other NUP98 FOs (25,54–57), it still displays incomplete penetrance and likely requires additional genetic alterations to drive leukemogenesis (Figure 1F). Nevertheless, these models represent valuable tools for future studies on the role of NUP98 FOs in various hematopoietic stem and progenitor populations and for functional genomic approaches. In this report, we demonstrate the utility of these models to uncover molecular dependencies in *NUP98-*r leukemia, such as MOZ and HBO1.

Pharmacologic inhibition of MOZ/HBO1 leads to myeloid differentiation and decreased global acetylation of H3K14 and H3K23 residues in *NUP98*-r cells. The HAT inhibitor PF9363 is specific for MOZ at low nanomolar concentrations, which is consistent with a global reduction in H3K23ac at these drug doses (Figure 4B). With higher nanomolar concentrations of PF9363, such as the 500 nM dose used in our in vitro experiments and the 3 mg/kg dose in our in vivo studies, it is possible that the therapeutic effect of PF9363 is also influenced by inhibition of HBO1. Consistent with this, *Moz, Hbo1, and Brpf1* were identified as dependencies in our in vivo CRISPR screen and/or CRISPR competition assays (Figure 3B, E). Furthermore, MOZ, HBO1, and BRPF1 were identified as interacting partners of NUP98 FO in our RIME studies (Figure 2B). While we cannot separate the impact of MOZ and HBO1 inhibition in many of the pharmacologic experiments reported here, the increase in CD11b expression observed at concentrations as low as 20 nM PF9363 in *NUP98*-r cells (Figure 4F-H) suggests that inhibition of MOZ alone is sufficient to induce myeloid differentiation ex vivo. Furthermore, we observed consistent decreases in H3K23ac after PF9363 treatment in vivo, while H3K14ac was more variable between animals (SFig 8N), suggesting that loss of H3K23ac is the primary consequence of PF9363 treatment at these concentrations.

Mechanistically, our RNAseq and ChIPseq studies suggest that treatment with PF9363 may displace FO from chromatin, analogous to results from our previous work with the Menin inhibitor VTP-50469 in *NUP98-*r cells (24). This was unexpected given that PF9363 is thought to inhibit the acetyltransferase domain of MOZ rather than disrupt protein-protein interactions (which is the proposed mechanism of action for Menin inhibitors). Consistent with our chromatin studies, a recent report in HEK293 cells expressing MOZ::TIF2 or CALM::AF10 demonstrated that the small molecule MYST HAT inhibitor WM-1119 can displace other FO from chromatin (59,60). Our results demonstrate that KMT2A and BRPF1 are also lost from chromatin after treatment with PF9363 monotherapy. We propose that NUP98 FOs serve as the anchor for the transcriptional complexes present at target gene loci; however, additional experiments using targeted degradation or other approaches are needed to elucidate how these complexes are maintained on chromatin at key pro-leukemogenic genes. Similar to what we have previously reported for Menin inhibitor Revumenib (24), we find that treatment with PF9363 and the combination of PF9363 and SNDX-5613 results in displacement of NUP98 FO at a subset of transcriptionally sensitive genes. At some loci, including *Meis1*, we observe a loss of chromatin binding as well as decrease in gene expression after treatment with the PF9363/SNDX-5613 combination. This finding nominates a small number of critical pro-leukemogenic genes (with shared RNA and chromatin changes) as the key mediators of drug response.

Finally, we show that MOZ/HBO1 inhibition with PF9363 is therapeutically effective in many *NUP98*-r models, with comparable efficacy to the recently FDA-approved Menin inhibitor SNDX-5613/Revumenib. Moreover, treatment with PF9363 can overcome Menin inhibitor resistance, suggesting that these agents may have distinct mechanisms for targeting key gene expression programs. While Menin inhibitors have demonstrated remarkable promise, reports of therapeutic resistance in patients enrolled on clinical trials (61) underscores the need for rational combination treatment. In our in vivo studies, PF9363 can synergize with SNDX-5613 to promote myeloid differentiation and prolong survival. This combination strategy may be readily translated as an early phase clinical trial of a MYST HAT inhibitor (NCT04606446; PF-07248144) in heavily pretreated breast cancer patients and demonstrated safety and tolerability (49). Given these results and our findings, MOZ/HBO1 inhibitors should be considered for further clinical testing as single agents and/or as a means of augmenting responses to Menin inhibition in cases of incomplete response or resistance in *NUP98*-r leukemia.

## MAIN METHODS

### Isolation and transduction of hematopoietic stem and progenitor cells (HSPCs)

Whole bone marrow was harvested from the forelimbs and/or hindlimbs of 6- to 12-week-old C57Bl/6 mice (RRID: IMSR_JAX:000664), and the EasySep Mouse Hematopoietic Progenitor Cell Isolation Kit (StemCell Technologies) was used according to the manufacturer’s instructions to select lineage negative hematopoietic stem and progenitor cells (lin-HSPCs). Cells were resuspended in IMDM (Gibco) containing 20% FBS (Hyclone), 1X penicillin-streptomycin-glutamine (Gibco), 50 ng/mL SCF (Peprotech), 40 ng/mL Flt3 (Peprotech), 30 ng/mL IL6 (Peprotech), 20 ng/mL IL3 (Peprotech), and 10 ng/mL IL7 (Peprotech) containing Lentiboost A or B (Sirion Biotech) at 1:100. Cells were then seeded in retronectin-treated plates pre-loaded with virus and spun at 800 x *g* and 4°C for 90 minutes. Cells were cultured at 37°C for 2-3 days and collected using cell dissociation buffer (Gibco). Transduced cells (mCherry-, Ametrine-, and/or GFP-expressing) were isolated using fluorescence-activated single cell sorting (FACS).

### Colony forming unit assay

Lineage negative HSPCs were plated in Methocult containing myeloid and erythroid growth factors (M3434, StemCell Technologies) and replated each week for 4 weeks, as described previously (18). The Nikon Eclipse TS100 microscope was used to acquire colony images.

### Generation of conditional knock in mouse model

*Nup98::Kdm5a* and *Nup98::Nsd1* conditional mouse models were created by Biocytogen. In the *Nup98::Kdm5a* model a loxP site was inserted upstream of *Nup98* exon 13. Downstream of this exon were inserted an FRT-flanked Neo-resistance positive-selection cassette, an mLoxP sequence, a reverse-orientation cDNA minigene containing *Nup98* exon 13, *Kdm5a* exon 27-HA, and poly-A tail, a LoxP sequence, and an mLoxP sequence, in that order. A similar design was used for *Nup98::Nsd1* mice with *Nup98* exon 12 and *Nsd1* exon 6-HA. Genotyping was performed on DNA extracted from tail or ear snips either using Transnetyx or PCR with KAPA2G Fast Genotyping Mix (Roche). Primer sequences are listed in Supplementary Table S1. To express *Nup98::Kdm5a* or *Nup98::Nsd1* specifically in the hematopoietic lineage, heterozygous fusion mice were crossed with *Vav*-Cre mice (62).

### Transplantation

Two days after isolating lineage negative HSPCs from *Nup98::Kdm5a;Vav*-Cre mice, *Nup98::Nsd1;Vav*-Cre mice, and wildtype *Vav-*Cre littermates, 200,000 HSPCs per recipient mouse were counted and resuspended in Phosphate Buffered Saline (PBS). 500,000 mature, CD117-negative bone marrow cells were isolated from a wildtype CD45.1 mouse using CD117 MicroBeads (Miltenyi Biotec) according to manufacturer’s recommendations. HSPCs and mature cells were combined in a total volume of 200 μL PBS per recipient. Cells were administered via tail vein injection into lethally irradiated (2 x 550 rad doses administered at least 3 hours apart using a cesium irradiator) CD45.1 primary recipients.

Secondary and tertiary recipients received 600,000 live spleen cells from mice of the previous passage, which were administered in 200 μL PBS via tail vein injection.

### CRISPR/Cas9 screening and analysis

Lineage negative HSPCs were isolated as above from Rosa26^Cas9^ knock in mice (Jackson Laboratory Cat No: 026179, RRID:IMSR_JAX:026179) and transduced to express NUP98::KDM5A and a guide RNA library targeting genes involved in epigenetic modulation (2022 sgRNAs from 337 epigenetic genes and 200 non-targeting sgRNAs) (40). The NUP98::KDM5A was expressed using a Gateway-compatible lentivirus vector (MND-mPGK-mCherry), and the guide RNA library was in a retroviral backbone with guide expression driven by the U6 promoter and an Ametrine reporter (41) and was prepared in PlatE cells. 1 million unsorted cells were injected per lethally irradiated (2 x 550 rad doses administered at least 3 hours apart using a cesium irradiator) CD45.2 primary mouse recipients.

FASTQ read files obtained after sequencing were demultiplexed using Hi-Seq analysis software (Illumina) and processed using MAGeCK (v.0.5.9.4) software (63). Raw count tables were generated using the MAGeCK count command by matching the guide sequence of the epigenetic library. Read counts for sgRNAs were normalized against median read counts across all samples for each screening. Volcano plots and log_2_ fold change of individual sgRNAs of enriched or depleted perturbations in indicated comparisons were depicted by MAGeCKFlute R package (v1.12.0). Two-tailed Fisher’s exact test was used to examine whether a Gene Ontology (GO) gene set was significantly (*P*L<L0.05) enriched among positive and negative regulators after genetic perturbation (significantly enriched or depleted (|log2(FC)|L> 0.5 and *P*L<L0.05) genes in BM tumor vs. spleen T cells and spleen tumor vs. spleen T cells).

### Co-culture assay

Non-tissue-culture-treated, 48-well plates were coated with RetroNectin (Takara) overnight at 4 deg C or at room temperature for 2 hours. Wells were washed with PBS, 750 μL gRNA virus (prepared in PlatE cells, gRNA sequences in Table S6) was added to each well, and plates were spun at 3000 rpm for at least 2 hours at 4°C. Wells were washed and cells were added with 1:100 Lentiboost A or B and spun for 90 minutes at 800*g* for 90 minutes. After 2-3 days of culturing in IMDM + 20% FBS + PSG + 10 ng/mL IL3, FACS was used to isolate GFP+ or Ametrine+ cells, which were cultured up to an additional 2 days in the same media after sorting. GFP+ and Ametrine+ cells were then counted and mixed in a ratio of approximately 1:1. Flow cytometry was used to determine the exact proportion of GFP+ and Ametrine+ cells at this timepoint and twice weekly for 3 weeks. At each analysis timepoint, cells were counted and replated at equal density.

### Rapid immunoprecipitation mass spectrometry of endogenous proteins (RIME)

HEK293T cells were transiently transfected to express empty vector, HA-NUP98, or HA-NUP98 FO and crosslinked using PBS with 1% formaldehyde (Sigma) after 48 hours. Nuclei were extracted and lysed, and sonication was performed using the Covaris E220. Immunoprecipitation was performed with HA antibody (Table S2) bound to protein G beads. Following washes with RIPA and ammonium hydrogen carbonate, peptides were purified with C18 spin columns prior to mass spectrometry analysis as previously described (64,65). Analysis was perfomed using the qPLEXanalyzer package (65). Please refer to **Supplemental Information** for additional details.

### Colocalization

HEK293T cells were seeded on poly-L-lysine-coated 15 mm coverslips of #1.5 thickness and transfected with GFP-tagged NUP98::LNP1 in the CL20 backbone as described previously (18). Twenty four hours after transfection, medium was removed and cells were washed with PBS and fixed with cold 100% methanol for 5 minutes at room temperature and 30 minutes at -20°C. Cells were then washed 3 x 5 minutes in PBS and coverslips were transferred to a humid chamber. Blocking was performed using 10% donkey serum for 1 hour at room temperature, then primary antibodies (Table S2) were added at 1:500 in 10% donkey serum and incubated overnight at 4°C. The next day, cells were washed 3 x 5 minutes in PBS + 0.1% Triton-X and incubated with secondary antibody (Table S2) at 1:300 in 5% donkey serum for 45 minutes. Three additional washes with PBS + 0.1% Triton-X were performed, each for 5 minutes. Cells were counterstained with 1 μM DAPI (Miltenyi Biotec) for 1-2 minutes and washed again 2 x 5 minutes with PBS + 0.1% Triton-X. Coverslips were mounted using anti-fade Prolong Diamond (ThermoFisher Scientific). Confocal images were acquired using the Marianas microscope. Individual nuclei were identified using Cellpose in 2D-mode, then combined across z-stacks by identifying which labels shared the most pixels in adjacent z-stacks. Puncta were segmented and quantified using the PunctaTools pipeline (18,66).

### RNA sequencing and analysis

For in vitro RNA-seq experiments on transformed NUP98::HOXA9 FO cells, cells were treated with DMSO, SNDX-5613, PF9363 or a combination of both agents, as indicated in figure legends. For in vivo RNA-seq experiments, PDX cells were harvested from the bone marrow of vehicle or PF9363 treated mice 54 days after transplantation, and human cells were enriched from the bone marrow of mice using the mouse cell depletion kit (Miltenyi Biotec # 130-104-694). RNA was extracted using the RNeasy kit (Qiagen). RNA (1 µg) was used to make Illumina compatible 3’-end libraries using QuantSeq 3’mRNA kit (Lexogen, Catalog numbers E7490L and E7530L). RNA-seq libraries were then sequenced on NextSeq550 (Illumina) to obtain 20-25 million unique sequencing reads. To perform gene-set enrichment analysis (GSEA), all the genes were ranked according to the fold-change and significance from differential analysis. GSEA was performed using mSigDB C2 genes.

### Global histone PTM mass spectrometry

Five million *Nup98::Kdm5a*;*Vav*-Cre;Cas9 HSPCs were cultured ex vivo in 5 mL media for 72 hours in the presence of DMSO or PF9363, then cells were washed, pelleted, and flash frozen. Samples were prepared and submitted for proteomic analysis by liquid-chromatography-mass spectrometry as described (67).

### Chromatin immunoprecipitation (ChIP) qPCR and sequencing (ChIPseq)

Cells were treated with either DMSO, SNDX-5613, PF9363 alone or in combination. At the indicated timepoint, cells were harvested and resuspended at 2.5x10^6^ cells/mL, then crosslinked using 1% methanol-free paraformaldehyde (ThermoFisher) for 10 minutes at room temperature. Glycine (25 mM) was used to quench the reaction. Cells were washed once with ice-cold PBS. Cytoplasm was stripped using 50 mM Tris-HCl pH 7.5, 140 mM NaCl, 1 mM EDTA, 10% Glycerol, 0.5% NP-40, and 0.25% Triton-X-100 for 10 min at 4°C. Nuclei were pelleted by centrifugation at 1400xg for 10 minutes. Nuclear pellets were then resuspended in 10 mM Tris-HCl pH 8.0, 200 mM NaCl, 1 mM EDTA and 0.5 mM EGTA, and incubated for 10 minutes at 4 degrees. Nuclei were pelleted at 1400xg for 10 minutes, then resuspended in 10 mM Tris-HCl pH 8.0, 100 mM NaCl, 1 mM EDTA, 0.5 mM EGTA, 0.1% sodium deoxycholate, 1% Triton-X-100, and 0.5% sodium lauroylsarcosine. Between 10-30 million cells (approximately 1-5 μg of chromatin) were used for each immunoprecipitation. Immunoprecipitation was performed using protein A (anti-Rabbit antibodies) or protein G (anti-mouse antibodies) magnetic beads (Dynal). Immunoprecipitated DNA fragments were eluted and de-crosslinked in 50 mM Tris-HCl pH 8.0, 10 mM EDTA, and 1% SDS, and quantified by TapeStation (Agilent) and Qubit (ThermoFisher). ChIP qPCRs used the SYBR green gene expression assays master mix (Applied Biosystems) and primers spanning the TSS and early exons of *Meis1* or the *Hoxa9* gene, as previously described (51). The ViiA 7 Real-Time PCR system (Applied Biosystems) was used with 384-well plates using SYBR green assays. Chromatin occupancy was calculated using the percent input method and normalized to the housekeeping gene *Gapdh*. Primer sequences appear in Table S1.

For ChIP-seq, 10-50 ng of chromatin was used to prepare Illumina-compatible libraries using a ThruPLEX DNA-Seq Kit (Takara) followed by sequencing using NextSeq550 (Illumina) to obtain 20-30 million unique sequencing paired-end reads per sample. For additional details on data analysis, please refer to **Supplemental Information**.

### Viability assay

Cells were treated with 0.1% DMSO, MOZ inhibitor (WM-1119 or PF9363), Menin inhibitor (VTP-50469 or SNDX-5613), or combination. Every 3 days, cells were replated with media containing fresh drug, as described previously (24). Viability was measured either by flow cytometry for DAPI as in (24) or using Cell Titer Glo (Promega) or resazurin. The Clariostar Plus plate reader was used to quantify Cell Titer Glo signal. Resazurin was prepared by dissolving 75 mg resazurin (Sigma), 12.5 mg methylene blue (Sigma), 164.5 mg potassium hexacyanoferrate (III), and 211 mg potassium hexacyoferrate (II) trihydrate (Sigma) to 500 mL sterile PBS (Gibco) and was stored in the dark at 4 deg C. The Biotek Synergy HTX plate reader was used to quantify resazurin signal using Gen5 3.12 software. Reading from the top of the plate, excitation fluorescence was collected at 530/25 nm and excitation wavelength was set to 590/35 nm with gain of 35. The signal collected from cells containing media only was used to remove background and normalize the data.

### Patient derived xenografts

NSG mice were transplanted with 0.5-1 million SJCML068699 (obtained from Prof. Cristina Mecucci (53)) or MSKG5191 PDX cells in 200 μL PBS via tail vein injection. Blood was collected retro-orbitally every 1-2 weeks to assess human/mouse chimerism by flow cytometry; antibodies are listed in Table S2. After 4 weeks, treatment with vehicles, PF9363, SNDX-5613 or combination was initiated (n=10/group). 3 mg/kg PF9363 was dissolved in 5 mL/kg 5% DMSO in 40% PEG300, 60% NaCl 0.9% and administered via intraperitoneal (IP) injection daily. 75 mg/kg SNDX-5613 was dissolved in 0.5% methylcellulose (type 400 cPs) in ultrapure water acidified with fumaric acid, at 10 mg/mL and was administered twice daily (with doses 6-8 hours apart) via oral gavage. Following treatment or at disease endpoint, mice were euthanized and blood, bone marrow, and spleen samples were collected and cryopreserved for analysis of leukemic burden.

For CPCT0021 or NTPL511 PDX experiments, female NSG mice (Jackson Laboratory Cat No: 005557, RRID:IMSR_JAX:005557) at 6-8 weeks of age housed in a pathogen-free animal facility at the Dana-Farber Cancer Institute. Mice were sublethally irradiated prior to tail vein intravenous injection of patient-derived xenograft cells. Engraftment of human cells in the peripheral blood was monitored every 1-2 weeks. Once engraftment was confirmed by the detection of human CD45+ cells in peripheral blood, treatment with vehicle, SNDX-5613 or PF9363 commenced. PF9363 3 mg/kg/dose was administered via intraperitoneal injection once daily for 5 days off, 2 days off, for a total of 30 days. SNDX-5613 was administered through ingestion of drug-formulated chow at the indicated dose. For the NTPL511 PDX survival experiment, animals were sacrificed when they developed signs of leukemia or >20% body weight loss, at which timepoint the bone marrow and spleen were harvested and analyzed using flow cytometry. For the CPCT0021 burden of disease experiment, all animals in the experiment were sacrificed at the same timepoint on Day 54, when mince in the control arm showed early signs of weight loss and rising human CD45+ percentage in the peripheral blood.

### Single cell RNA sequencing

MSKG5191 PDX cells were injected into NSG mice and allowed to engraft to ∼30% (based on flow cytometry for human CD45) in the peripheral blood. Mice were treated for 4 weeks with vehicle, SNDX-5613, PF9363, or combination as described above. Mice were humanely euthanized, and spleen samples were cryopreserved. Human cells were isolated using the EasySep mouse/human chimera isolation kit (StemCell Technologies), according to manufacturer’s instructions. From each treatment group or vehicle viable PDX cells from the spleen of 2 different mice were equally pooled and pooled samples were washed 3 times with 1X PBS (calcium and magnesium free) containing 0.04% weight/volume BSA (Thermo Fisher Scientific, catalogue number AM2616) and automatedly counted by the Countess 3 Automated Cell Counter (Thermo Fisher Scientific). From each pooled sample, we calculated the volume of cells to load to have a desired recovery target of 8,000-10,000 cells and loaded on Chromium Next GEM Chip K (10X Genomics; PN-2000182) with Master Mix from Chromium Next GEM Single Cell 5’ Reagent Kits v2 (Dual Index) (10X Genomics; PN-1000263), Single Cell VDJ 5’ Gel Beads (10X Genomics; PN-1000264) and Partitioning Oil (10X Genomics; PN-2000190) following standard manufacturer’s protocols and as previously described (68,69). Illumina-ready dual index libraries were sequenced at the recommended depth and aiming at approximately 35,000-50,000 reads/cell on a NovaSeqX according to manufacturer’s recommendations.

### Single cell RNAseq analysis

Single-cell RNA-seq data were aligned and quantified using the Cell Ranger (v7.0.1) pipeline (http://www.10xgenomics.com) against the human genome GRCh38 (refdata-gex-GRCh38-2020-A). When the mitochondrial content for a cell was more than 1.5 times the interquartile range (IQR) greater than the 75^th^ percentile of mitochondrial content for the total cell population, the cell was excluded. Clusters of cells were identified using Seurat (5.1.0, https://satijalab.org/seurat) using Uniform Manifold Approximation and Projection (UMAP). Cell type annotation was defined by using the reference bone marrow previously described (69,70). Loupe Browser (version 8) was used to visualize the data.

### Assay for transposase-accessible chromatin (ATAC)-seq

MSKG5191 PDX cells were injected into NSG mice and allowed to engraft to ∼30% (based on flow cytometry for human CD45) in the peripheral blood. Mice were treated for 5 days with vehicle, SNDX-5613, PF9363, or combination as described above. Mice were humanely euthanized, and human cells were isolated from their spleens using the EasySep mouse/human chimera isolation kit (StemCell Technologies), according to manufacturer’s instructions. Spleen samples were cryopreserved, then viably thawed and washed with PBS. 50,000 cells were subjected to cell lysis, nuclei extraction, tagmentation and library preparation as described (71,72). Lysis and transposition were adjusted according to the Omni-ATAC protocol (73). Libraries were prepared as described in (71) with double-sided bead purification and were sequenced using a NovaSeq targeting 50 million paired-end reads per sample. For additional details on data analysis, please refer to **Supplemental Information**.

### Data accessibility

Data has been deposited and is accessible using GEO Accession Numbers GSE283096 and GSE283532.

## Acknowledgements

We thank the Hartwell Center for Biotechnology, the Vector Core Lab, the Flow Cytometry and Cell Sorting Shared Resource, the Genome Sequencing Facility, the Center for Applied Bioinformatics, the Comprehensive Pathology Core, the Center for Advanced Genome Editing, the Cell and Tissue Imaging Center and the Animal Resource Center, the Preclinical Therapeutics Program, and the Analytical Technologies Center-Preclinical Formulation Team (ATC-PFT) of St. Jude Children’s Research Hospital.

We thank Dana-Farber/Harvard Cancer Center in Boston, MA, for the use of the Specialized Histopathology Core, which provided histology and immunohistochemistry service. Dana-Farber/Harvard Cancer Center is supported in part by an NCI Cancer Center Support Grant # NIH 5 P30 CA06516. We also thank Dr. Jerry McGeehan from Syndax Pharmaceuticals.

