## Supplementary Methods and Figures for "MOZ and HBO1 Histone Acetyltransferase Complexes Are Molecular Dependencies and Therapeutic Targets in *NUP98*-Rearranged Acute Myeloid Leukemia"

SUPPLEMENTARY MATERIAL

**Supplemental Methods**

*Cell Culture*

HEK293T cells were obtained from ATCC (RRID:CVCL_1926), authenticated by STR-profiling (PowerPlex Fusion), and maintained in DMEM with 10% fetal bovine serum (FBS, Hyclone) and 1X penicillin-strepomycin-glutamine (Gibco). PlatE cells were cultured in DMEM with + 10% fetal bovine serum (FBS, Hyclone) and 1X penicillin-strepomycin-glutamine (Gibco). PlatE media contained 1 ug/mL puromycin and 10 ug/mL blasticidin except during virus preparation.

Mouse hematopoietic stem and progenitor cells expressing NUP98 fusion oncoproteins (via viral overexpression or conditional knock in) were maintained in culture with IMDM (Gibco) containing 20% FBS (Hyclone), 1X penicillin-streptomycin-glutamine (Gibco), and 10 ng/mL IL3 (Peprotech).

The mouse NUP98::HOXA9, NUP98::KDM5A, and NUP98::NSD1 *Wt1*-/- transformed cell lines were maintained in IMDM with 15% FBS, 1% penicillin/streptomycin (Gibco), 6 mg/mL mIL3, 10 mg/mL mIL6, and 10 mg/mL mSCF (StemCell Technologies). All cells were maintained at 37 º C and 5% CO2, except for PlatE cells during virus preparation.

*Virus Preparation*

As described previously (1), retroviruses were prepared in HEK293T cells using FuGENE HD transfection reagent (Promega, E2311) and high-titer lentiviruses were. PlatE cells were used to prepare gRNA viruses. Briefly, 800,000 cells per well were plated in 6-well dishes. The following day, media was changed 2 hours before transfection. 3 ug of the gRNA plasmid and 1 ug of pCL-Eco were added to 250 uL of Opti-MEM (Gibco) along with 10 uL of Transit-293 (Mirus Bio). The mixture was added to cells after 20-minute incubation at room temperature. The media was changed after 12-18 hours, and after 24 more hours the cells were incubated at 32ºC for the final 32 hours until the viral supernatant was harvested with 0.45 uM filtering. Viruses were used the same day or flash-frozen and stored at -80 ºC until needed.

*Inhibitors*

SNDX-5613 (cat# HY-136175, MedChemExpress), VTP-50469 (cat# HY-114162, MedChemExpress) PF9363 (cat# HY-132283, MedChemExpress), and WM-1119 (cat# HY-102058, MedChemExpress) were dissolved in DMSO at 10 mM and diluted in media to indicated concentrations for in vitro experiments.

*Animals*

Experiments involving C57Bl/6 mice, conditional knock in *Nup98::Kdm5a* and *Nup98::Nsd1* mice, and PDXs SJCML068699 and MSKG5191 were performed at St. Jude Children's Research Hospital. All mice were treated ethically under Animal Care and Use Committee (IACUC)-approved protocol number 584-100506-10/23.

Experiments involving PDXs NTPL511 and CPCT0021 were performed with the approval of the Dana-Farber IACUC.

*Complete blood counts and stained blood smears*

Complete blood counts and stained blood smears were performed by the St. Jude Comprehensive Pathology Core using the Genesis analyzer and Aerospray Hematology Pro Slide Stainer/Cytocentrifuge, respectively.

*Immunohistochemistry*

IHC for conditional knock in mouse tumors was performed on formalin-fixed paraffin-embedded tissues sectioned at 4µm. All assay steps for CD3, RUNX1, GATA1, MPO, and GlyA, including deparaffinization, rehydration, and epitope retrieval, were performed on the Ventana Discovery Ultra autostainer with Ventana Reaction Buffer (Ventana Medical Systems, 950-300) rinses between steps. Antibody binding was detected using the OmniMap Rabbit Detection kit (Roche, 760-4311) for 16 minutes (CD3 first labeled with a rabbit anti-goat secondary antibody, and RUNX1 with a rabbit anti-mouse antibody), followed by ChromoMap DAB (Roche, 760-159) for 10 minutes. All assay steps for Ly6B, PAX5, and B220 were performed on the Bond Max with Bond wash buffer (Leica, AR9590) rinses between steps. Slides were incubated with the primary antibody; antibody binding was detected using the anti-rabbit Bond Polymer Refine Detection kit (Leica, DS9800), with Ly6B and B220 first labeled with a rabbit anti-rat secondary antibody. All assay steps for CD41 were performed on the Biocare intelliPATH with Biocare TBS wash buffer (cat # TWB954M, Biocare) rinses between steps. Slides were incubated with the primary antibody followed by the rabbit-on-rodent HRP polymer (Biocare Medical, RMR622) and then DAB (ThermoFisher, TA-125-HDX).

Immunohistochemistry (IHC) for CPCT0021 PDX bone marrow cells was performed after fixation in 10% formalin and embedding in paraffin for sectioning. Sections were stained with Haemotoxylin and Eosin (H&E) to visualize hematopoietic cells. IHC was performed on the Leica Bond III automated staining platform. Antibody CD45 was run at 1:50 dilution with citrate antigen retrieval using the Leica Biosystems Refine Detection Kit. VWF antibody was used at 1:1000 dilution with citrate antigen retrieval using the Leica Biosystems Refine Red Detection kit. See Table S2 for additional details on specific antibodies used for IHC.

*Flow cytometry*

For immunophenotypic analysis of cells from colony forming unit assays (cryopreserved in FBS with 10% DMSO after each week of growth in Methocult, 0.5-1 uL of each antibody (see Table S2) was used for 20-30 minutes in the dark at 4°C. After staining, cells were washed again in PBS + 2% FBS (Hyclone) and 1 μL of DAPI staining solution (Miltenyi Biotec) was added immediately prior to analysis. Immunophenotypic analysis of peripheral blood, bone marrow, and spleen samples was performed similarly immediately after isolation of cells from mice. Antibodies are listed in Table S2. All data were collected on a BD Fortessa flow cytometer and analyzed using FlowJo (v10.7.1).

*Rapid immunoprecipitation mass spectrometry of endogenous proteins (RIME)*

For RIME experiments, 8 million HEK293T cells were plated per 15-cm dish. The following day, cells were transfected with 30 ug per plate of plasmid (empty vector, HA-NUP98, or HA-NUP98 fusion) in the CCLMPC backbone. 120 uL of FuGENE HD transfection reagent (Promega, E2311) was added to 1.5 mL of Opti-MEM (Gibco) for 5 minutes, then plasmid was added. After a 20-30 minute incubation, the mixture was added to cells dropwise. Media was removed 48 hours after transfection, and cells were scraped off the plates into 15 mL conical tubes using PBS (Gibco) with 1% formaldehyde (Sigma). Cells were rotated for 10 minutes at room temperature, then crosslinking was quenched using 0.5 mL of 2.5 M glycine for 5 minutes. Cells were pelleted by centrifugation at 1500 rpm and 4°C for 3 minutes, then washed twice in PBS + 1X protease inhibitors (Roche) with centrifugation at 8000 rpm and 4 °C for 3 minutes. Pellets were then flash frozen on dry ice and stored at -80 deg C or used immediately for lysis and sonication.

One mL of lysis buffer 1 (LB1; 50 mM HEPES-KOH pH7.5, 140 mM NaCl, 1 mM EDTA, 10% glycerol, 0.5% Igepal CA-630, 0.25% Triton X-100) was used to resuspend each pellet, and samples were rotated at 4°C for 10 minutes. The supernatants were removed after centrifugation at 2000 x g and 4°C for 5 minutes, and each pellet was resuspended in 1 mL lysis buffer 2 (LB2; 10 mM Tris-HCl pH 8.0, 200 mM NaCl, 1 mM EDTA, 0.5 mM EGTA). Samples were rotated at 4°C for 5 minutes, then centrifuged for 5 minutes at 2000 x g and 4°C. Next each pellet was resuspended in lysis buffer 3 (LB3; 10 mM Tris-HCl, 100 mM NaCl, 1 mM EDTA, 0.5 mM EGTA, 0.1% Na-deoxycholate, 0.5% N-lauroylsarcosine). Each pellet was resuspended in 2 mL of LB3, 1 mL of which was transferred to 2 separate 1 mL millitubes with AFA fiber (Covaris). Samples were sonicated for 17 minutes using the Covaris E220 with 5% duty cycle, 200 cycles per burst, 140 Watts, and water level 8. Sonicated material was transferred to 2 mL Low-Bind tubes, and Triton-X was added to a final concentration of 1%. Samples were centrifuged for 10 minutes at 20,000*g* and 4°C, then supernatants were transferred to new tubes. Samples were diluted to 1.5 mL per tube in LB3 + 1% Triton-X. A 10 uL aliquot of sonicated material was kept for reverse crosslinking and assessment of fragment sizes on a 2% DNA gel; the desired fragment size was 200-300 bp. 50 uL of Protein G beads per sample were previously prepared by washing 3 times with 0.5% BSA in PBS then incubating with 5 ug HA antibody (Abcam ab9110) for 1 hour at room temperature or 4°C overnight. After 3 additional washes with 0.5% BSA in PBS, beads were resuspended in 200 uL LB3 + 1% Triton-X per sample and added to individual tubes. Antibody bound beads and samples were rotated overnight at 4°C, then washed 10 times with 1 mL RIPA buffer (50 mM HEPES pH 7.6, 1 mM EDTA, 0.7% Na-deoxycholate, 1% Igepal CA-630, and 0.5 M LiCl) and twice with 500 uL of 100 mM ammonium hydrogen carbonate. Finally, beads were flash frozen and submitted for tryptic digestion and mass.

Peptides were purified with C18 spin columns prior to mass spectrometry analysis as previously described (6,7). Peptides were analyzed with an UltiMate 3000 UHPLC system coupled with the Q-Exactive HF (Thermo Scientific) mass spectrometer. The full scans were performed in the Orbitrap in the range of 400 to 1600 m/z at 60k resolution. For MS2 scans a resolution of 30k was selected with 2.0 Th isolation window, HCD collision energy 28% and dynamic exclusion 30 seconds. MS2 spectra were searched against a human UniProt database containing over 20,000 entries including custom FOs protein sequences (Table S3) in Proteome Discoverer 2.4. The SequestHT node included the following parameters: Precursor Mass Tolerance 20 ppm, Fragment Mass Tolerance 0.02 Da, number of maximum missed cleavages sites 2 and Dynamic Modifications were Oxidation of M (+15.995Da) and Deamidation of N/Q (+0.984Da). The confidence level for peptide identifications was estimated with the Percolator node using target-decoy database search. Label-free quantification was performed with the Minora Feature Detector node. The consensus workflow included the Feature mapper and the Precursor Ion Quantifier node using intensity for precursor quantification. Only unique peptides were used for downstream statistical analysis. Missing values in the empty vector samples and in the various bait protein peptides were replaced with zeros. Peptide intensities were normalized using a median scaling normalization, and the KNN method (8) was used to impute missing values. Peptides with greater than 50% missing values were filtered-out. Protein level quantification was obtained by summing the normalized and imputed peptide intensities. Using the qPLEXanalyzer package (7), a statistical analysis to identify differentially regulated proteins was conducted. Multiple hypothesis testing correction of p-values was applied using the Benjamini-Hochberg method (9) to control the false discovery rate (FDR). Preranked Gene Set Enrichment Analysis (GSEA) was carried out using GSEA (version 4) (10).

*Immunoprecipitation and western blot*

Twenty million adherent HEK293T cells were seeded in a 10 cm plate and transfected with 10 µg HA-NUP98::KDM5A, HA-NUP98::HOXA9 or HA-NUP98::LNP1 plasmid or empty vector control plasmid using FuGENE HD transfection reagent (Promega, E2311) as described above for RIME experiments. The cells were collected 72 hours post transfection by centrifugation at 300*g* for 3 minutes. The cell pellet was washed with cold PBS and stored at -80°C until further use.

Cell pellets were lysed in 500 ml of lysis buffer containing M-PERTM mammalian protein extraction reagent (Thermo Scientific, 78501) and 1x cOmplete protease inhibitor cocktail (Millipore Sigma, 11836153001), at 4°C. The lysate was cleared by centrifugation at 18000*g* at 4°C for 5 minutes. Total protein estimation was performed on the clarified lysate using BCA method. An aliquot was saved for loading controls. Approximately 2mg/ml of the clarified lysate was incubated with 50ml of HA antibody agarose (Pierce, 26181) for 1hour at 4°C on a rotisserie. The agarose gel was precipitated by centrifugation at 250*g* for 2 minutes and washed twice with 500 ml of wash buffer containing 50mM Tris-HCl pH 7.5 and 150 mM NaCl. Samples were prepared for SDS-PAGE by adding 25 µl 1x Protein loading dye to the immunoprecipitated agarose samples and by boiling the samples for 2 minutes at 95°C. Loading controls were prepared similarly.

Samples were run a 4-12% gradient SDS-PAGE at 120V constant for 1.5 hours and the proteins were transferred to a nitrocellulose membrane using semi-dry method (Invitrogen-iBLOT2). The nitrocellulose membranes were blocked with 5% milk at room temperature for 1 hour. The blots were incubated overnight at 4°C on a rocker with primary antibody, then incubated with at room temperature for 1 hour on a rocker. The blots were washed 3 times with TBST, developed with ECL reagent (BioRad, 1705060) and imaged on a BioRad ChemiDoc imager.

*Whole genome sequencing*

For conditional knock in mouse cells transplanted into CD45.1 recipients, FACS was used to isolate CD45.2+ spleen cells after staining with CD45.2-FITC antibody (see Table S2). For conditional knock in mice with spontaneous leukemia, unfractionated spleen cells were used. For each, DNA was isolated using the Qiagen AllPrep kit. Mouse genomic libraries were generated using the SureSelectXT kit specific for the Illumina HiSeq instrument (G9611B, Agilent Technologies). Whole whole genome sequencing were performed on the Illumina HiSeq with 150 bp reads by the St. Jude Hartwell Center for Biotechnology. Tumor/pseudo-control matched variant calling was performed, with the normal sample from a different mouse used as pseudo-control. An ensemble strategy with optimized filtering (SNVs/Indels called by at least 2 callers) was employed to identify potential somatic coding mutations (SNVs/indels) using multiple callers, including Mutect2 (v4.1.2.0)(11), SomaticSniper (v1.0.5.0)(12,13), VarScan2 (v2.4.3)(14), MuSE (v1.0rc)(15), and Strelka2 (v2.9.10)(16). To exclude mouse germline SNPs, variants observed in either mouse dbsnp142 from NCBI or mouse EVA SNP databases from the European Variation Archive (EVA) at the European Bioinformatics Institute (EBI) were excluded. Variant annotation was performed using Annovar(17). These consensus calls, particularly for coding mutations that potentially disrupt protein coding sequences, further underwent manual assessment and evaluation of read depth, mapping quality, and strand bias, in order to eliminate additional artifacts.

*CD34+ HSPC culture and treatment*

Cord blood-derived CD34+ HSPCs were transduced to express HA-NUP98::KDM5A as previously described and were cultured in StemSpan-XF (StemCell Technologies) containing 1% penicillin-sprepomycin, 50 μg/mL each human IL-6, Flt-3 ligand, SCF, and TPO (Peprotech), as well as 10 mM each StemReginin-1 (StemCell Technologies) and UM729 (Stem Cell Technologies). Cells were cultured in DMSO or 100 nM PF9363, with cells counted every 3-5 days, washed with PBS, and replated in media with fresh DMSO/inhibitor for a total of 19 days. Statistical analysis was performed with a generalized linear mixed effect model using glmer (lmer) function (lmer package, v1.1-35.1) and emmeans function (emmeans package, v1.5.0), and with unpaired two-sided Student’s t-test with equal variances (pairs function).

*ChIP-seq data analysis*

Raw Illumina sequencer output was converted to FASTQ format using bcl2fastq (v2.20.0.422). Reads (paired-end 37-mers) were trimmed for quality using trimmomatic (v0.36; minimum trimmed length 34bp), aligned to the mouse genome (Gencode M24/mm10) using STAR (v2.7.5a), sorted and duplicates marked/removed with picard pipeline tools (v2.9.4). Final ‘‘deduped’’ .BAM files were indexed using SAMtools (v1.95). Total signal was assessed around TSS regions using the sitepro tool (v0.6.6) from the Cis-regulatory Element Annotation System (CEAS) package, based on signal from WIG files generated using IGVtools (v2.3.98). Peaks were called using MACS2, and input samples were used as controls for peak calling. TSS region intervals were defined (-1kb to +3kb) using sitepro from the CEAS suite. Positive signal was defined for genes that displayed a 2-fold enrichment above the input sample, with at least > 200 reads within the aforementioned TSS window. To compare DMSO samples with each drug condition, a single representative TSS region was identified for each gene, based on the site with the highest average signal in DMSO control samples, and a ratio of DMSO to drug treatment was calculated for each gene. Data visualizations were produced using IGVtools (TDF signal pileups), deepTools (3.5.5.) and ngs.plot (pileup heatmaps).

*ATAC-seq data analysis*

To perform statistical test between experimental groups, the number of fragments were counted for each reference peak using the intersect command from pybedtools (v0.8.1) (18,19). Next, the number of raw reads mapping per peak was converted to FPKM unit (Fragments Per Kilo base per Million mapped reads), and TMM (trimmed mean of M-values) using edgeR (20). Finally, the limma-voom approach (21,22) was used to assess the significance of the differential peak binding / accessibility. For motif enrichment analysis, Homer software (v4.10) was used for comparison to known motifs (23).

**Supplemental Results**

*Characterization of conditional knock in Nup98::Kdm5a and Nup98::Nsd1 mice*

As show in Figure 1E, we observed increased spleen weight in 3-month-old *Nup98::Kdm5a;Vav*-Cre mice as compared to wildtype littermates. No differences in mature cell composition were noted between groups in the bone marrow or spleen (SFig 1B). *Nup98::Kdm5a;Vav*-Cre mice had significantly more lin- cells than control mice, with a decrease in the percentage of cKit+ cells within this population and no other significant differences in composition (SFig 1C). There were no significant changes noted for *Nup98::Nsd1;Vav*-Cre mice, consistent with their weaker phenotype in the CFU assay (Figure 1D, SFig 1B-C).

After transplantation of HSPCs from *Nup98::Kdm5a*;*Vav*-Cre, *Nup98::Nsd1;Vav*-Cre*,* or wildtype *Vav*-Cre mice into lethally irradiated congenic recipients, we observed more rapid engraftment of *Nup98::Kdm5a;Vav*-Cre cells compared to wildtype *Vav-*Cre cells, with significantly more donor (CD45.2+) cells in the peripheral blood after 4 weeks (SFig 1D). The resulting *Nup98::Kdm5a;Vav-*Cre leukemias were characterized by splenomegaly, often accompanied by high white blood cell counts and anemia (SFig 1E-G). Bone marrow and spleen cells from sick animals variably expressed Gr1 and B220 (but not CD3 or CD19) while showing stronger positivity for CD11b, MPO, and Ly6B (SFig 1H-I). Transplantation of spleen cells from leukemic mice into secondary and tertiary recipients resulted in similar disease phenotypes within 4 weeks (Figure 1F, SFig 1E-H). Whole genome sequencing of leukemic cells revealed no recurrent high quality, coding mutations between primary recipients (Table S4).

Monitoring of conditional knock in mice expressing NUP98::KDM5A revealed a survival difference between *Nup98::Kdm5a;Vav*-Cre and *Vav-*Cre littermates (SFig 2A). From a cohort of *Nup98::Kdm5a;Vav*-Cre mice monitored over approximately two years, 25 out of 30 (83%) mice become moribund, of which 5 (20% of moribund mice, 17% of total) were confirmed to have myeloid disease. In contrast, 5 out of 25 (20%) of *Vav*-Cre mice monitored over the same time became moribund, and we did not identify any cases of myeloid disease in this group (SFig 2A). While disease in the latter group of 5 *Nup98::Kdm5a;Vav-*Cre animals universally displayed expression of myeloid markers (CD11b, Gr1, MPO by immunophenotyping and/or immunohistochemistry), mice exhibited varying splenomegaly and white blood cell counts (SFig 2B-E). We performed serial transplantation experiments using spleen cells from three *Nup98::Kdm5a;Vav-*Cre mice with AML confirmed by necropsy. In two of these three cases (67%), secondary recipients became moribund with median latencies of 27 and 48 days (SFig 2F). Recipients from the third animal showed no signs of disease after approximately four months. We performed genomic analysis of three *Nup98::Kdm5a;Vav-*Cre mice that spontaneously developed myeloid disease, which revealed only one recurrent SNV in *Kmt2a* (Table S5). Interestingly, the same mutation (T1311I) was reported previously as a recurrent mutation in hematologic malignancies from a *Dnmt3a* knockout mouse model (24).

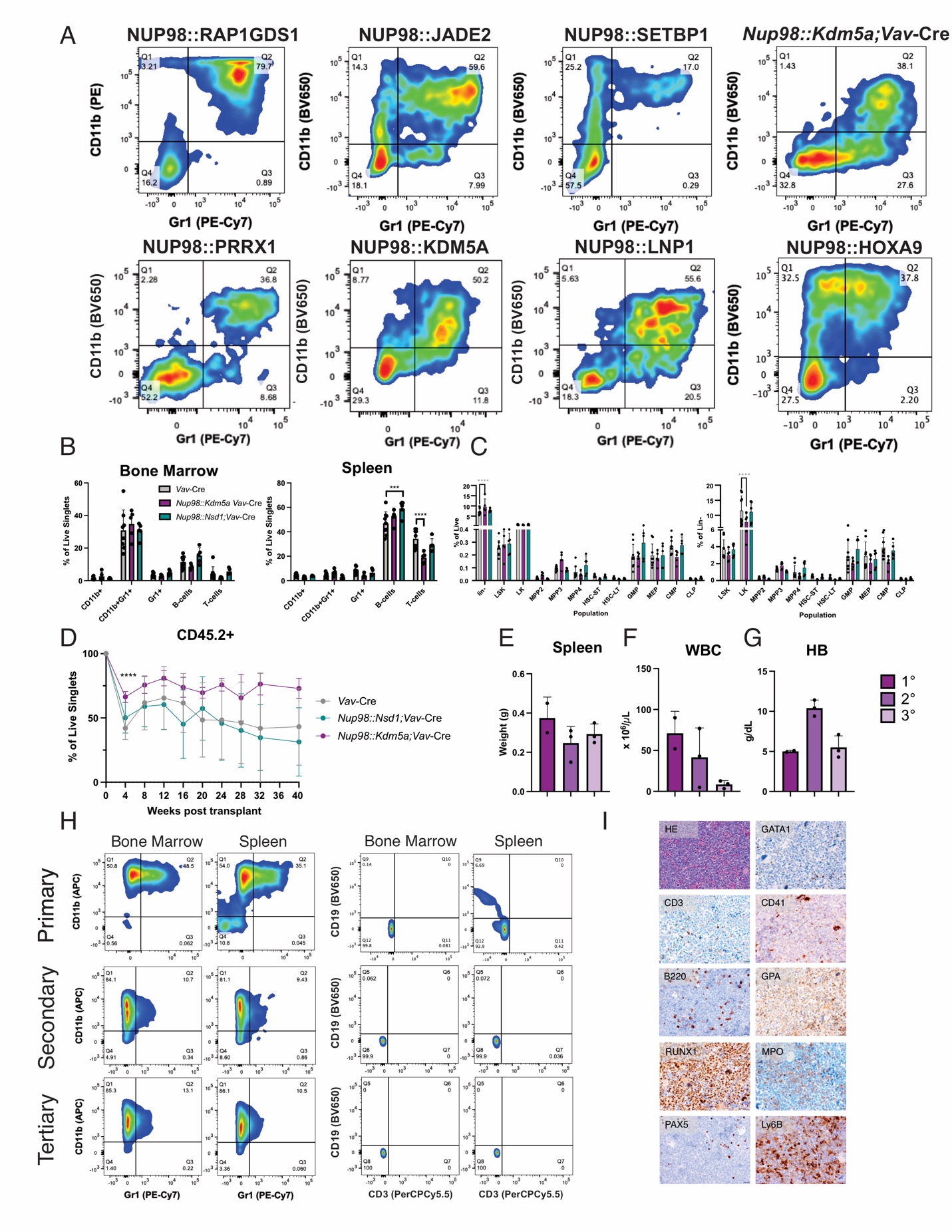

**Supplemental Figure 1** *(Related to Figure 1)****.*** Some *NUP98* fusions drive cell transformation and leukemogenesis.

A) Immunophenotyping of lin- HSPCs expressing NUP98 FOs after two (NUP98::KDM5A, LNP1, PRRX1) or three (NUP98::HOXA9) weeks of growth in methylcellulose containing myeloid and erythroid growth factors. Data shown are from the mEGFP-positive live singlet population in a representative experiment. NUP98::HOXA9, KDM5A, LNP1, and PRRX1 results have been published previously(1) and are included for comparison. Immunophenotyping was performed on bone marrow and spleen samples from conditional knock in *Nup98::Kdm5a;Vav-*Cre or *Nup98::Nsd1;*Vav-Cre mice or wildtype *Vav-*Cre littermates, and quantification of B) mature and C) stem/progenitor populations is shown. D) Lin- HSPCs from *Nup98::Kdm5a;Vav-*Cre mice, *Nup98::Nsd1;*Vav-Cre mice, or wildtype *Vav-*Cre littermates (CD45.2) were transplanted into lethally irradiated congenic (CD45.1) recipients. Engraftment was measured over time. E) Spleen weight, F) white blood cell count, and G) hemoglobin after the development of leukemia in recipients of *Nup98::Kdm5a;Vav-*Cre lin- HSPCs. H) Representative immunophenotyping of bone marrow and spleen of mice after primary, secondary, or tertiary transplant of *Nup98::Kdm5a*;*Vav*-Cre lin- HSPCs. I) Representative immunohistochemistry of spleen from primary recipient of *Nup98::Kdm5a*;*Vav*-Cre lin- HSPCs after leukemia development. 40X magnification, scale bar 20 μm.

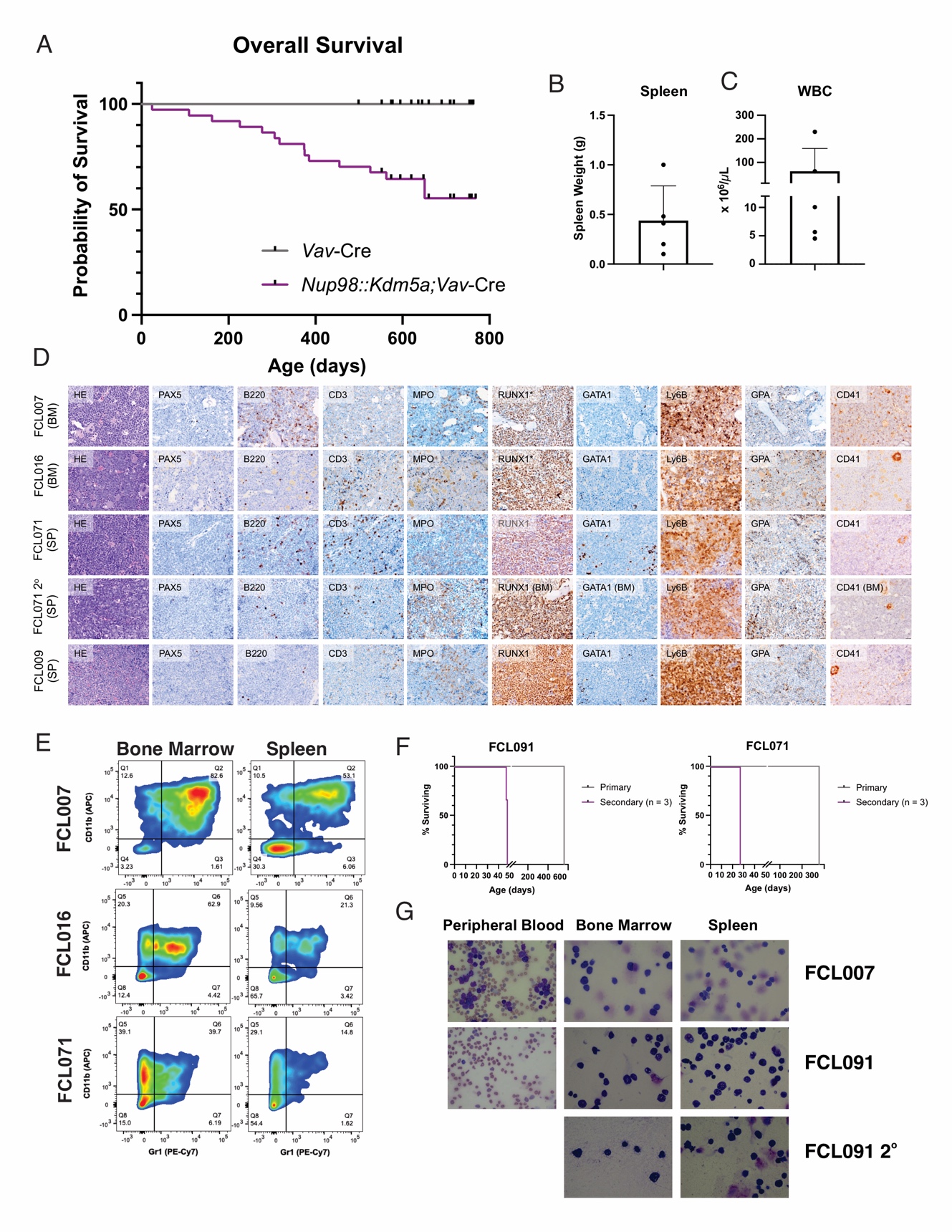

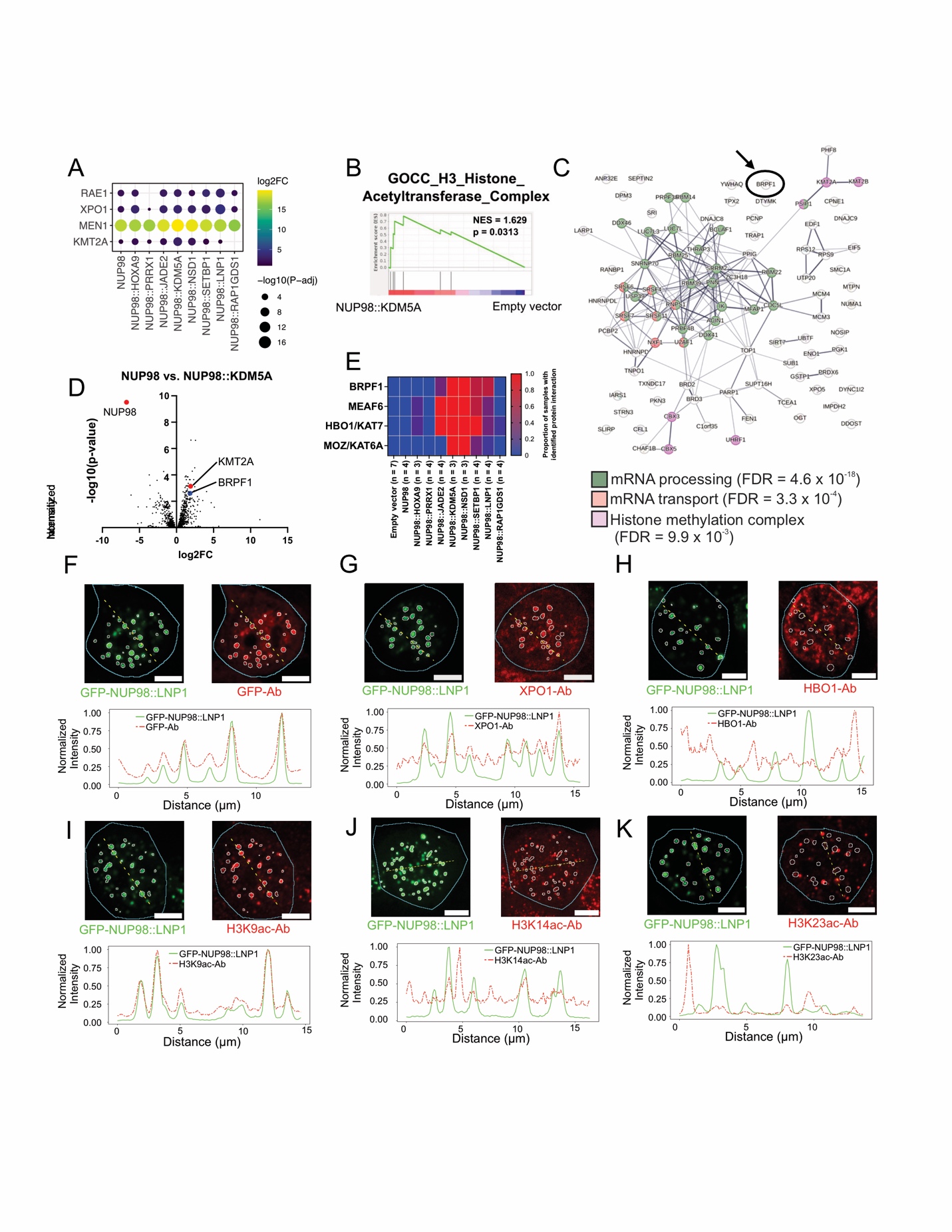

**Supplemental Figure 2** *(Related to Figure 1)****.*** Nup98::Kdm5a;Vav-Cre mice spontaneously develop myeloid disease.

A) Overall survival curve for *Nup98::Kdm5a;Vav-*Cre mice and wildtype littermates. B) Spleen weight and C) white blood cell count after the development of myeloid disease in *Nup98::Kdm5a;Vav-*Cre mice. D) Representative immunohistochemistry of bone marrow or spleen samples from *Nup98::Kdm5a*;*Vav*-Cre mice after development of myeloid disease. 40X magnification, scale bar 20 μm. E) Representative immunophenotyping of bone marrow and spleen of *Nup98::Kdm5a;Vav-*Cre mice after disease development. F) Survival curve for *Nup98::Kdm5a;Vav-*Cre mice that spontaneously developed leukemia and secondary recipients of their spleen cells. G) Representative images of stained blood smear and bone marrow or spleen cytospin for spontaneous *Nup98::Kdm5a;Vav-*Cre leukemias. 40X magnification.

**Supplemental Figure 3** *(Related to Figure 2).* NUP98 FOs interact with histone acetyltransferase complex members.

A) Bubble plot showing log2 fold change and p-value for interaction of NUP98 and NUP98 FOs with RAE1, XPO1, MEN1, and KMT2A. B) Volcano plot for enriched proteins in isolated by label-free quantification of RIME in transiently transfected HEK293T cells expressing HA-NUP98::KDM5A compared to HA-NUP98. Data with missing values in the empty vector were imputed. The average log2 fold-change of 3-4 biological replicates was used for the plot, and interaction of interest are labeled. C) STRING analysis of enriched proteins (log2FC > 1, adjusted p-value < 0.025) from RIME in transiently transfected HEK293T cells expressing HA-NUP98::KDM5A as compared to HA-NUP98. D) Gene set enrichment analysis of NUP98::KDM5A FO interactors pre-ranked based on significance versus empty vector samples. E) Heatmap showing proportion of samples in which HAT complex members were identified by RIME for empty vector, NUP98, or NUP98 FO. HEK293T cells were transiently transfected with GFP-tagged NUP98::LNP1, fixed with methanol, and stained for F) GFP, G), XPO1, H) HBO1, I) H3K9ac, J) H3K14ac, or K) H3K23ac primary antibodies and RRX-conjugated secondary antibodies. Images were acquired by confocal microscopy. Top panels show example cells with GFP-NUP98::LNP1 in green and antibody signal in red. GFP condensate segmentation is shown with white outlines and cell nuclear boundaries are shown with cyan outlines.  The yellow dotted line shows a path where normalized intensities are plotted in the lower panel. Scale bar, 5 μM.

**Supplemental =** *(Related to Figure 3).* Histone acetyltransferase complex members are molecular dependencies in *NUP98*-rearranged cells.

A) Gating strategy to isolate FO+gRNA+ (mCherry+Ametrine+) myeloid (leukemic) BM and spleen cells from mice after in vivo epigenetic CRISPR/Cas9 screen. T-cells were also sorted as a control. B) Log2 fold change of individual gRNAs for commonly depleted gRNAs (*Bmi1*, *Prdm12*, *Brwd3*, *Brpf1*) in BM and spleen NUP98::KDM5A leukemia samples after in vivo epigenetic CRISPR/Cas9 screen. C) Relative fitness of *Nup98::Kdm5a;Vav*-Cre;Cas9 HSPCs transduced with gRNAs targeting HAT complex members (marked with Ametrine) and placed in competitive co-culture with similar cells transduced with non-targeting control gRNA (marked with GFP). Second replicate, consistent with data in Figure 3E. D) Relative fitness of NUP98::LNP1-expressing Cas9+ HSPCs transduced with gRNAs targeting HAT complex members (marked with Ametrine) and placed in competitive co-culture with similar cells transduced with non-targeting control gRNA (marked with GFP). Representative data are shown.

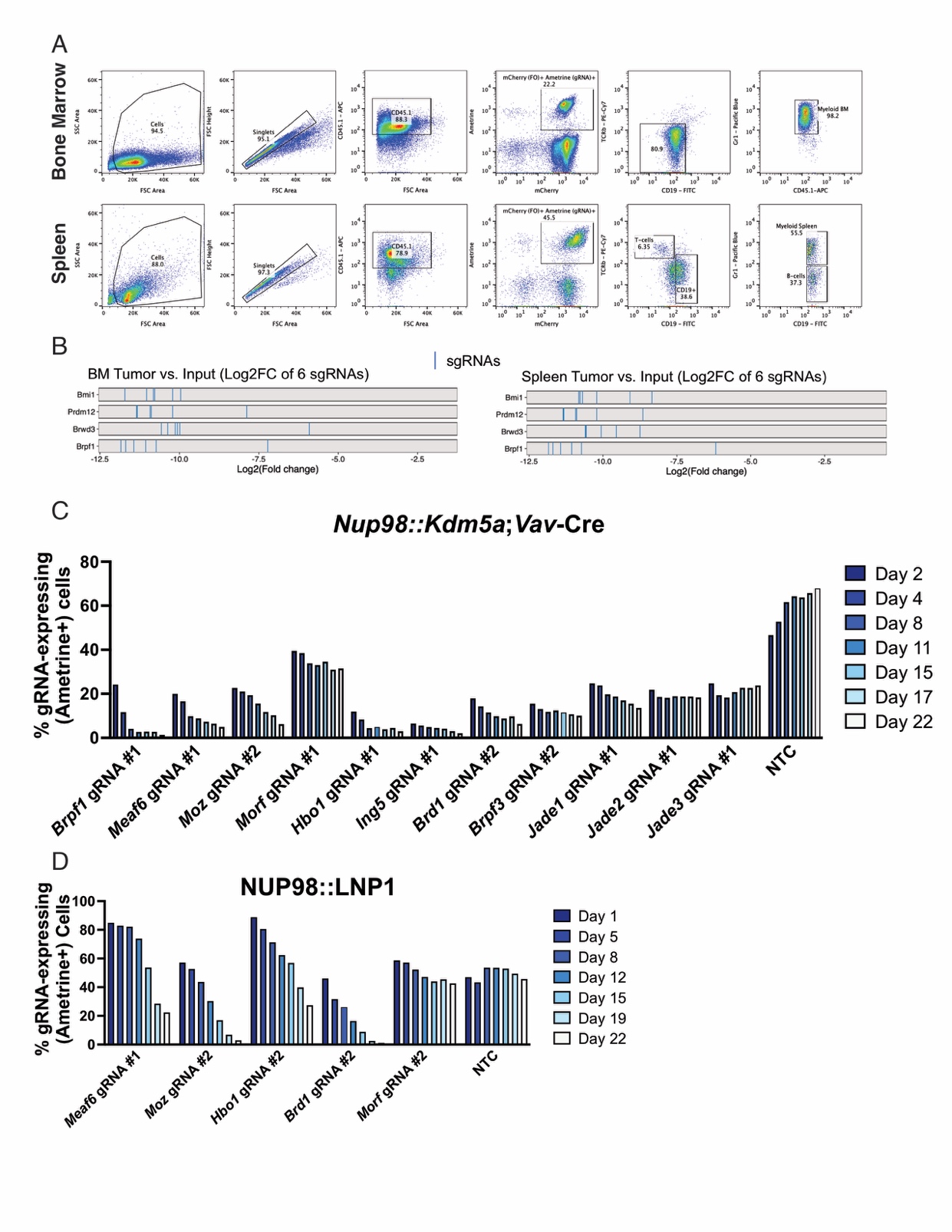

**Supplemental Figure 4** *(Related to Figure 3).* Histone acetyltransferase complex members are molecular dependencies in *NUP98*-rearranged cells.

A) Gating strategy to isolate FO+gRNA+ (mCherry+Ametrine+) myeloid (leukemic) BM and spleen cells from mice after in vivo epigenetic CRISPR/Cas9 screen. T-cells were also sorted as a control. B) Log2 fold change of individual gRNAs for commonly depleted gRNAs (*Bmi1*, *Prdm12*, *Brwd3*, *Brpf1*) in BM and spleen NUP98::KDM5A leukemia samples after in vivo epigenetic CRISPR/Cas9 screen. C) Relative fitness of *Nup98::Kdm5a;Vav*-Cre;Cas9 HSPCs transduced with gRNAs targeting HAT complex members (marked with Ametrine) and placed in competitive co-culture with similar cells transduced with non-targeting control gRNA (marked with GFP). Second replicate, consistent with data in Figure 3E. D) Relative fitness of NUP98::LNP1-expressing Cas9+ HSPCs transduced with gRNAs targeting HAT complex members (marked with Ametrine) and placed in competitive co-culture with similar cells transduced with non-targeting control gRNA (marked with GFP). Representative data are shown.

**Supplemental Figure 5** *(Related to Figure 4).* *NUP98*-r cells respond to MOZ inhibition in vitro.

A) Volcano plots of histone posttranslational modifications identified by mass spectrometry after treatment of *Nup98::Kdm5a;Vav*-Cre-Cas9 HSPCs for 72 hours with vehicle (DMSO) or varying concentrations of PF9363. B) PBMCs from two donors were treated for 72 hours with increasing concentrations of PF9363, and viability was measured using a resazurin cell viability assay. C) CD34+ cells were treated with vehicle (DMSO) or increasing concentrations of PF9363 in methylcellulose for 2 weeks, then colonies were counted and scored for morphology.

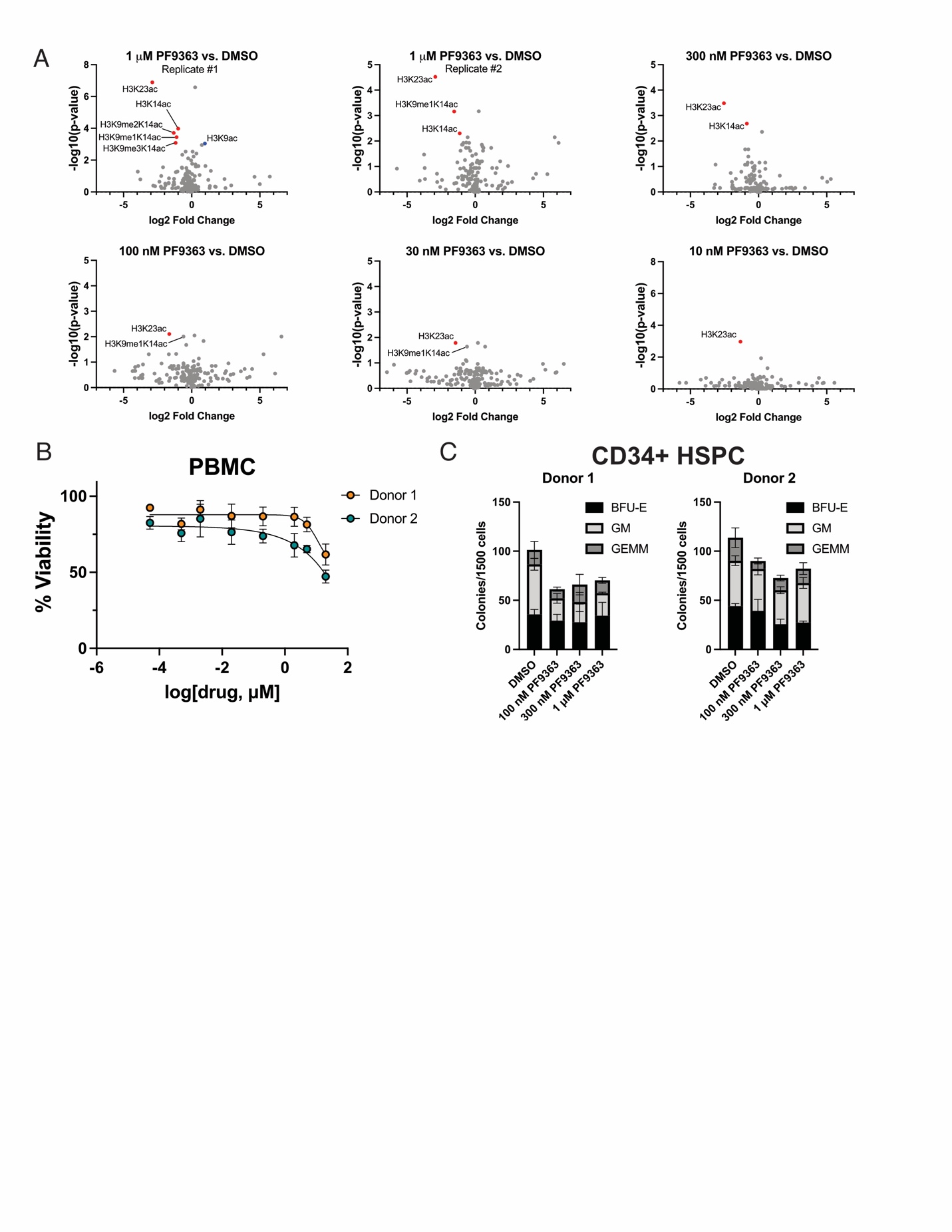

**Supplemental Figure 5** *(Related to Figure 4).* *NUP98*-r cells respond to MOZ inhibition in vitro.

A) Volcano plots of histone posttranslational modifications identified by mass spectrometry after treatment of *Nup98::Kdm5a;Vav*-Cre-Cas9 HSPCs for 72 hours with vehicle (DMSO) or varying concentrations of PF9363. B) PBMCs from two donors were treated for 72 hours with increasing concentrations of PF9363, and viability was measured using a resazurin cell viability assay. C) CD34+ cells were treated with vehicle (DMSO) or increasing concentrations of PF9363 in methylcellulose for 2 weeks, then colonies were counted and scored for morphology.

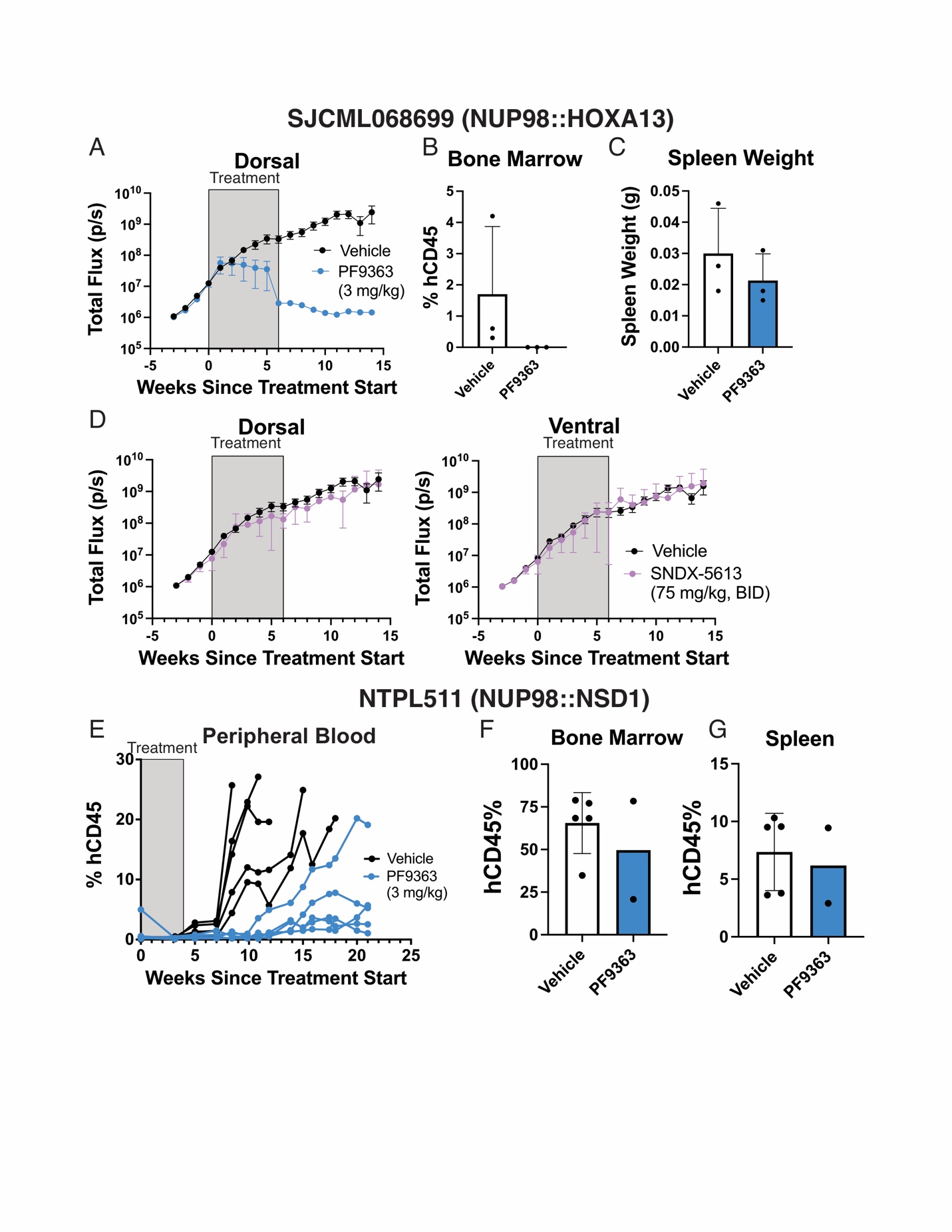

**Supplemental Figure 6** *(Related to Figure 5).* *NUP98*-r PDXs respond to MOZ inhibition in vivo.

A-C) Mice bearing luciferase-marked NUP98::HOXA13 PDX cells were treated with vehicle or PF9363 (3 mg/kg, 5 days on, 2 days off, IP) for 6 weeks. A) Total dorsal flux was monitored over time. Leukemia burden was measured at the end of the 6-week treatment by B) hCD45 in bone marrow and C) spleen weight. D) Leukemia burden was measured by IVIS in mice bearing luciferase-marked NUP98::HOXA13 PDX cells during treatment with vehicle or SNDX-5613 (75 mg/kg, 5 days on, 2 days off, BID) for 6 weeks. E) hCD45 was measured in the peripheral blood by flow cytometry for mice bearing NUP98::NSD1 PDX cells during treatment with vehicle or PF9363 (3 mg/kg, QD, IP) for 4 weeks. At the time of sacrifice, leukemia burden was measured by F) hCD45 in bone marrow and G) spleen weight.

**Supplemental Figure 6** *(Related to Figure 5).* *NUP98*-r PDXs respond to MOZ inhibition in vivo.

A-C) Mice bearing luciferase-marked NUP98::HOXA13 PDX cells were treated with vehicle or PF9363 (3 mg/kg, 5 days on, 2 days off, IP) for 6 weeks. A) Total dorsal flux was monitored over time. Leukemia burden was measured at the end of the 6-week treatment by B) hCD45 in bone marrow and C) spleen weight. D) Leukemia burden was measured by IVIS in mice bearing luciferase-marked NUP98::HOXA13 PDX cells during treatment with vehicle or SNDX-5613 (75 mg/kg, 5 days on, 2 days off, BID) for 6 weeks. E) hCD45 was measured in the peripheral blood by flow cytometry for mice bearing NUP98::NSD1 PDX cells during treatment with vehicle or PF9363 (3 mg/kg, QD, IP) for 4 weeks. At disease endpoint, leukemia burden was measured by F) hCD45 in bone marrow and G) spleen weight.

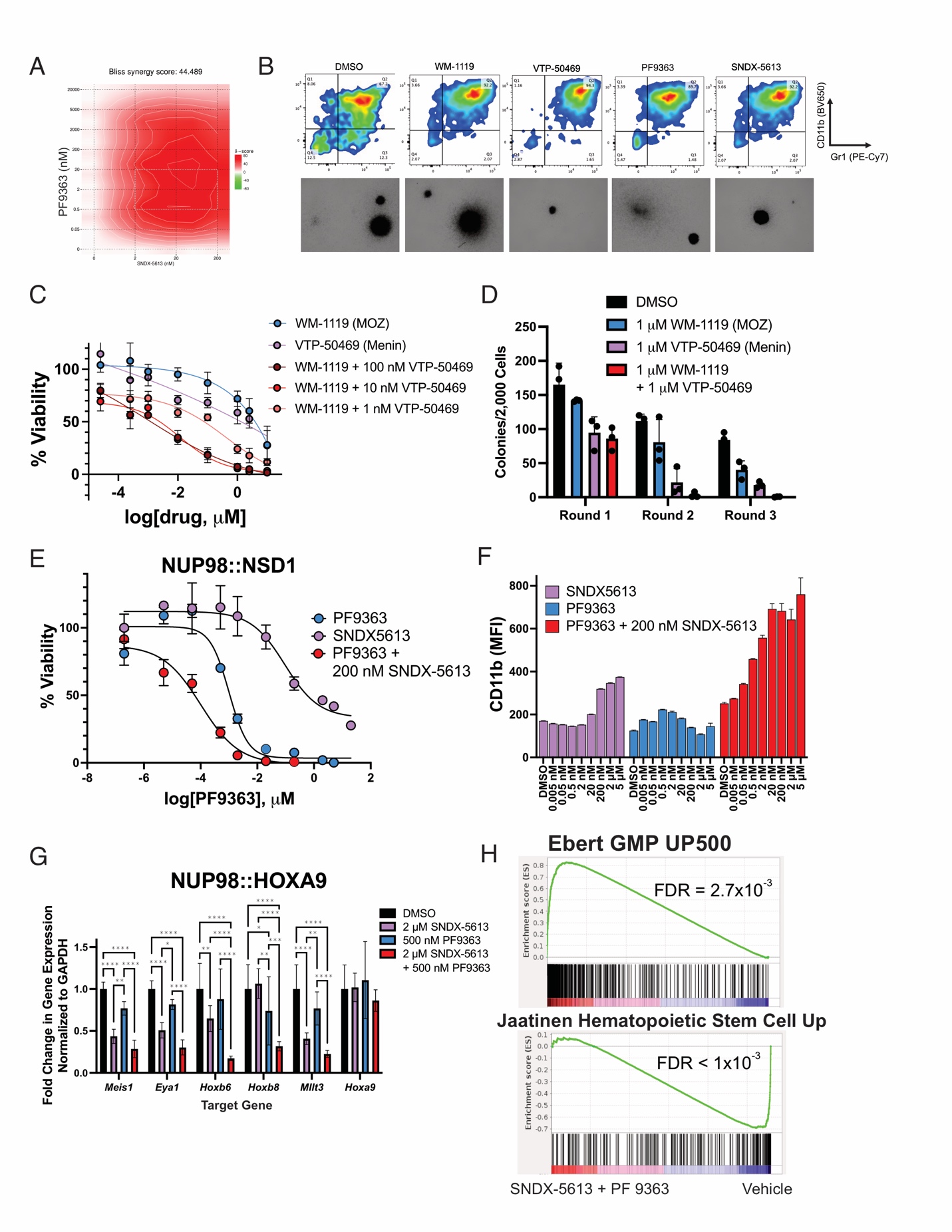

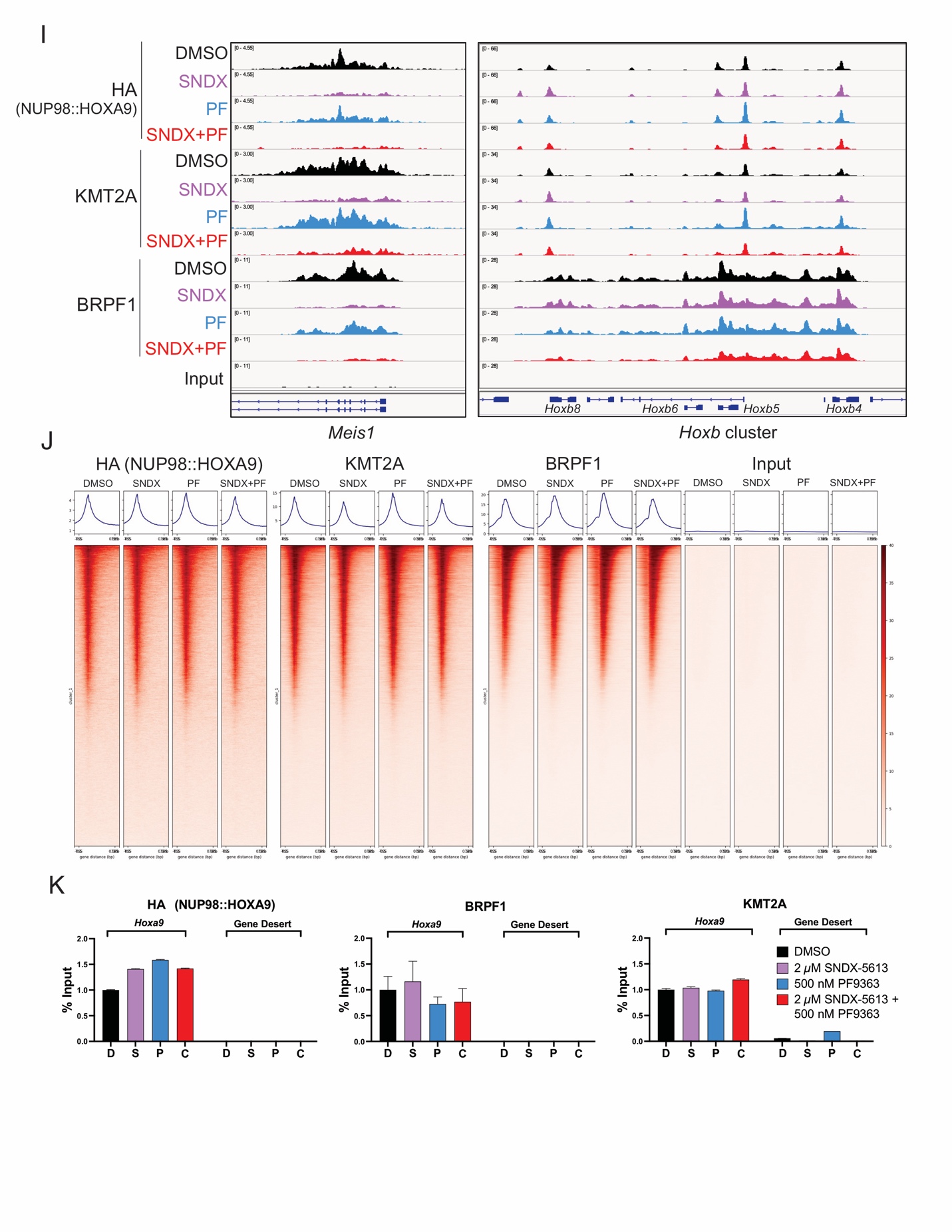

**Supplemental Figure 7** *(Related to Figure 6).* MOZ and Menin inhibition alter gene expression and remodel chromatin in *NUP98*-r cells.

A) Heat map showing Bliss synergy scores for *Nup98::Kdm5a;Vav*-Cre;Cas9 HSPCs treated with PF9363, SNDX-5613, or combination for 9 days. Plot was generated using data from Figure 6A using SynergyFinder (3). B) Immunophenotyping and images of *Nup98::Kdm5a;Vav*-Cre lin- HSPCs after three weeks of growth in methylcellulose containing myeloid and erythroid growth factors. Flow cytometry data shown are from the live singlet population in a representative experiment. C) *Nup98::Kdm5a;Vav*-Cre;Cas9 HSPCs were treated with WM-1119, VTP-50469, or combination for 9 days, and viability was measured relative to vehicle (DMSO)-treated controls using a resazurin cell viability assay. D) Colony forming unit assay for *Nup98::Kdm5a;Vav*-Cre lin- HSPCs treated with vehicle (DMSO), 1 μM WM-1119, 1 μM SNDX-5613, or combination and replated each week with fresh inhibitors. NUP98::NSD1 mouse leukemia cells were treated with PF9363, SNDX-5613, or combination for 10 days, and E) viability or F) CD11b expression was measured compared to vehicle (DMSO)-treated controls. G) qPCR results after treatment of NUP98::HOXA9 mouse leukemia cells for 72 hours with vehicle (DMSO), 2 uM SNDX-5613, 500 nM PF9363, or combination. Mean +/- SD by 2-way ANOVA and Tukey’s multiple comparison test, * denotes *P* > 0.05, ** denotes *P* < 0.01, *** denotes *P* < 0.001, and **** denotes *P* < 0.0001. Data shown are from a representative experiment. H) Gene set enrichment analysis for differentially expressed genes after 2 uM SNDX-5613 and 500 nM PF9363 treatment for 72 hours in NUP98::HOXA9 mouse leukemia cells, compared to vehicle (DMSO) treatment. Upregulation of Ebert GMP UP500 gene set (4) and dowregulation of Jaatinen Hematopoietic Stem Cell Up gene set (5) was observed after combination treatment compared to vehicle. I) IGV tracks and J) tornado plots for HA (NUP98::HOXA9), BRPF1, and KMT2A binding by ChIPseq in NUP98::HOXA9 mouse leukemia cells after treatment with vehicle (DMSO), 2 uM SNDX-5613, 500 nM PF9363, or combination for 72 hours. Data shown are from a representative experiment.K) ChIP-qPCR for HA (NUP98::HOXA9), BRPF1, and KMT2A at the *Hoxa* locus or gene desert in NUP98::HOXA9 mouse leukemia cells after treatment with vehicle (DMSO), 2 uM SNDX-5613, 500 nM PF9363, or combination for 72 hours.

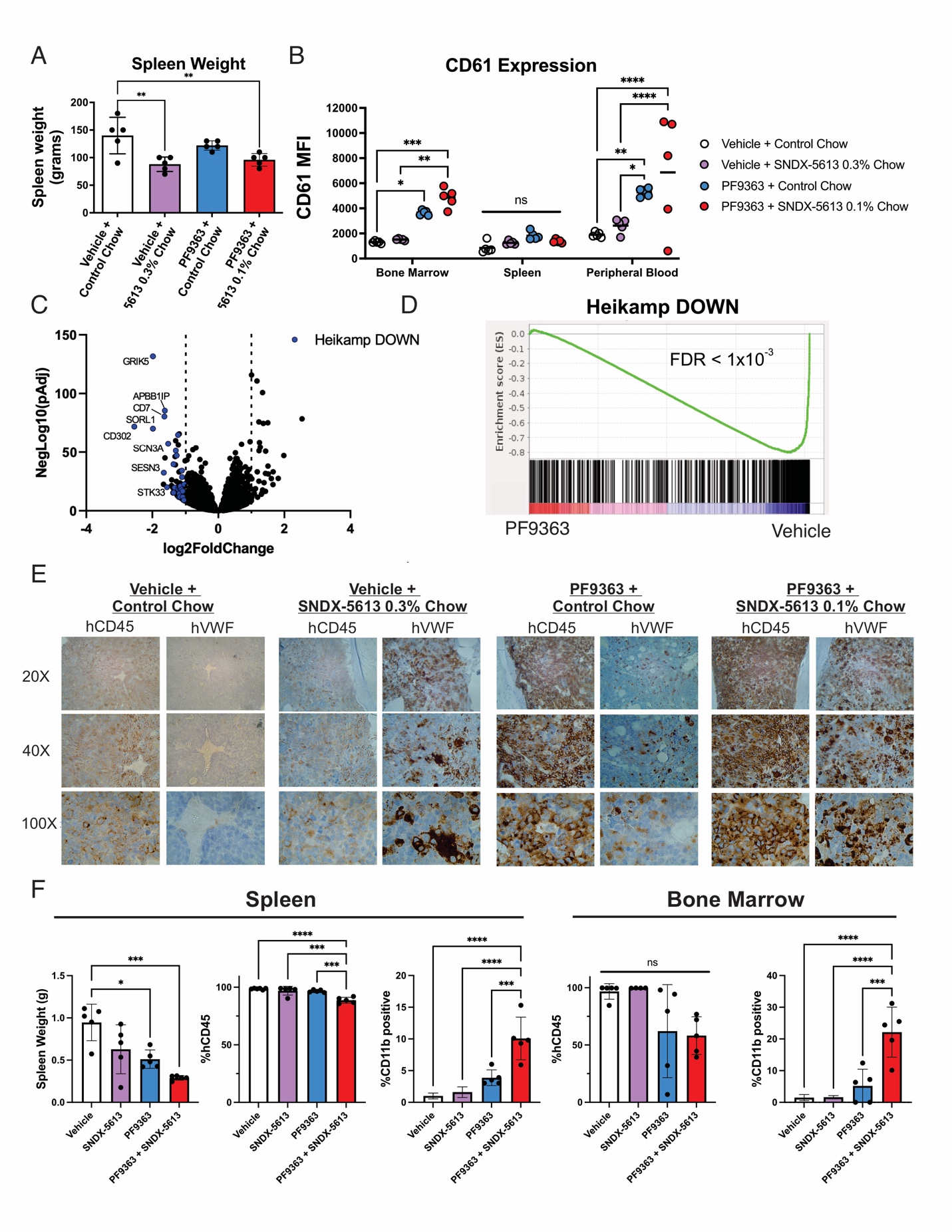

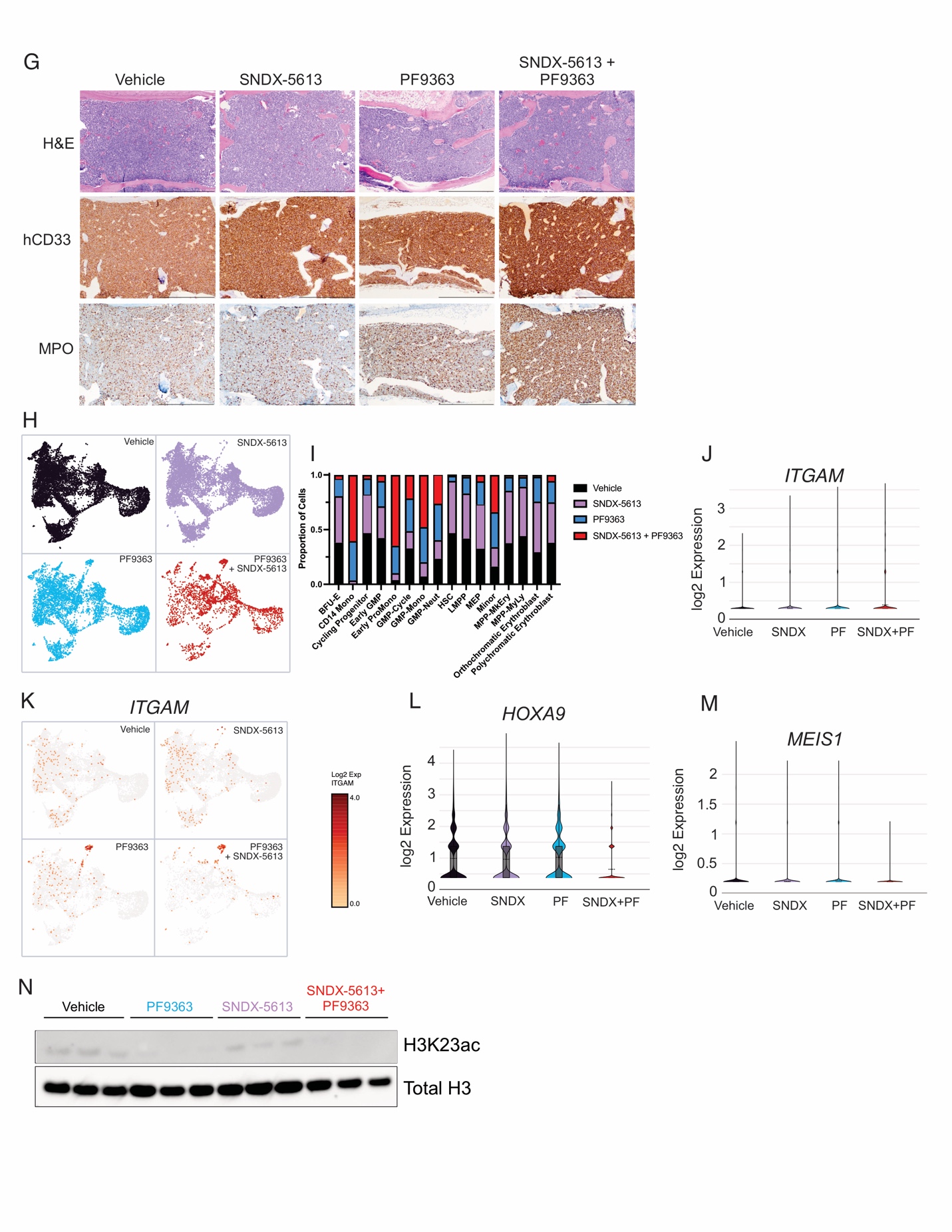

**Supplemental Figure 8** *(Related to Figure 7).* MOZ and Menin inhibition have combinatorial effects in vivo.

Mice bearing CPCT0021 (NUP98::KDM5A) PDX cells were treated with vehicles, PF9363 (3 mg/kg, QD, IP), SNDX-5613 (0.3% chow), or combination (3 mg/kg PF9363, QD, IP + 0.1% SNDX-5613 chow). A) Human CD61 expression was measured at the end of treatment in the bone marrow, spleen, and peripheral blood. C) Volcano plot showing differentially expressed genes in RNA-seq bone marrow samples from vehicle or PF9363-treated mice. D) Gene set enrichment analysis of downregulated genes in NUP98::KDM5A PDX cells treated with vehicle or PF9363.(2) E) Immunohistochemistry for human CD45 (hCD45) or VWF (hVWF) in bone marrow samples after 4-week treatment. 20X, 40X, and 100X magnification. F) Mice bearing MSKG5191 (NUP98::NSD1) PDX cells were treated for 4 weeks (5 days on, 2 days off) with vehicles, PF9363 (3 mg/kg, QD, IP) , SNDX-5613 (75 mg/kg, BID, PO), or combination, and bone marrow and spleen samples were harvested. Spleen weight was measured, and immunophenotyping was performed to determine the percentage of human CD45 and CD11b positive cells. G) Immunohistochemistry for hCD45 and CD33 was performed on sternum samples at the end of 4-week treatment. 10X magnification, scale bar 500 μm. H-M) Single cell RNA-seq was performed using the 10X Genomics 3’ technology in spleen samples from n = 2 mice per treatment arm (pooled samples) after 4 weeks of treatment. H) UMAP projection, separated by sample/treatment arm. I) Proportion of cells in each cluster from each sample/treatment arm. J) Violin plot and K) UMAP projection for *ITGAM* expression between treatment arms. Violin plot for expression of L) *HOXA9* and M) *MEIS1* in each sample/treatment arm. N) MSKG5191 PDX-bearing mice were treated for 5 days with vehicle, PF9363, SNDX-5613, or combination using the doses above (n = 3/group), then histones were isolated from human leukemic cells in the spleen. H3K23ac and total histone H3 expression were probed by western blot.
